# Genetic analysis of the X-linked Adrenoleukodystrophy *ABCD1 gene* in *Drosophila* uncovers a role in Peroxisomal dynamics

**DOI:** 10.1101/2024.09.23.614586

**Authors:** Joshua Manor, Sharayu V Jangam, Hyung-lok Chung, Pranjali Bhagwat, Jonathan Andrews, Hillary Chester, Shu Kondo, Saurabh Srivastav, Juan Botas, Ann B. Moser, Suzette M. Huguenin, Michael F Wangler

## Abstract

X-linked adrenoleukodystrophy (X-ALD) is a progressive neurodegenerative disorder caused by a loss-of-function (LOF) mutation in the *ATP-binding cassette subfamily D member 1* (*ABCD1)* gene, leading to the accumulation of very long-chain fatty acids (VLCFAs). This disorder exhibits striking heterogeneity; some male patients develop an early childhood neuroinflammatory demyelination disorder, while other patients, including adult males and most affected female carriers, experience a chronic progressive myelopathy. Adrenocortical failure is observed in almost all male patients, with age of onset varying sometimes being the first diagnostic finding. The gene underlying this spectrum of disease encodes an ATP-binding cassette (ABC) transporter that localizes to peroxisomes and facilitates VLCFA transport. X-ALD is considered a single peroxisomal component defect and does not play a direct role in peroxisome assembly. *Drosophila* models of other peroxisomal genes have provided mechanistic insight into some of the neurodegenerative mechanisms with reduced lifespan, retinal degeneration, and VLCFA accumulation. Here, we perform a genetic analysis of the fly ABCD1 ortholog *Abcd1* (CG2316). Knockdown or deficiency of *Abcd1* leads to VLCFA accumulation, salivary gland defects, locomotor impairment and retinal lipid abnormalities. Interestingly, there is also evidence of reduced peroxisomal numbers. Flies overexpressing the human cDNA for *ABCD1* display a wing crumpling phenotype characteristic of the *pex2* loss-of-function. Surprisingly, overexpression of human *ABCD1* appears to inhibit or overwhelm peroxisomal biogenesis to levels similar to null mutations in fly *pex2*, *pex16* and *pex3*. *Drosophila Abcd1* is therefore implicated in peroxisomal number, and overexpression of the human *ABCD1* gene acts a potent inhibitor of peroxisomal biogenesis in flies.

## INTRODUCTION

X-linked adrenoleukodystrophy (X-ALD) is a peroxisomal disorder resulting in the defective breakdown of very long chain fatty acids (VLCFA, C ≥ 22), affecting myelin and axons in the central nervous system (CNS) and the adrenal gland, culminating in a progressive neurometabolic disorder. X-ALD occurs due to mutations in the X-linked *ABCD1* gene, which encodes a peroxisomal transmembrane protein responsible for the transport of CoA-esters of VLCFA (CoA-VLCFA) into the peroxisome. X-ALD exhibits an estimated birth incidence of 1 in 17,000, rendering it the most common hereditary peroxisomal disorder.^1,2^ A defining feature of X-ALD is the systemic accumulation of saturated VLCFA, notably hexacosanoic acid (C26:0). A sensitive, fast, high-throughput assay for C26:0 LPC facilitated the addition of X-ALD to the RUSP newborn screening panel.^2^

X-ALD presents pronounced phenotypic heterogeneity, underscored by an absolute absence of genotype-phenotype correlation. The default phenotype of X-ALD is a slowly progressive spinal cord axonopathy, called adrenomyeloneuropathy (AMN). AMN is observed in at least of 40% males and upwards of 80% of females. Yet, 35-40% of affected males develop childhood cerebral ALD (ccALD), an early onset, rapidly progressing inflammatory cerebral demyelination, representing a more severe phenotype. Additionally, 20% of males will “transition” from an AMN phenotype to a later-onset cerebral X-ALD. Adrenal insufficiency is almost uniformly presented in male patients, either in conjunction with ccALD or AMN or, less frequently, as a distinct phenotype of ‘adrenal insufficiency only’.^3–5^ The pathology differentiating cerebral ALD (childhood or later onset) from AMN is starkly distinct: ccALD is characterized by demyelination, oligodendrocyte loss in the CNS, with activated microglia, astrocytes, and macrophage infiltration, whereas AMN typically exhibits minimal to no CNS reactive astrocytosis or lymphocytosis, but significant symmetric spinal cord atrophy, particularly in the lateral corticospinal, gracile, and spinocerebellar tracts, alongside distal axonopathy and myelin loss.^5^ The disparate phenotypes and hence management approaches for ccALD, AMN, and adrenal insufficiency only have spurred intense interest in identifying early neuroinflammation biomarkers for ccALD. Yet, the ability to determine a specific X-ALD phenotype by existing biomarkers or genetic variants remains unpredictable.

The gene *ABCD1* is located on Xq28, spanning 19.9 kbp and comprises 10 exons. It encodes for an ATP binding cassette subfamily D member 1 (ALDP, or ABCD1). ABCD1 dimer binds CoA-VLCFA to its transmembrane domain (TM, exons 1-2), and ATP to its cytosolic nucleotide binding domain (NBD, exons 6-9). The import is biphasic, initially involving the attachment of VLCFA-CoA to a thioesterase domain followed by an ATP-dependent substrate hydrolysis along with a conformational change inducing the opening of the TM towards the peroxisomal lumen.^6,7^ Currently, the X-ALD variant database (ALD info, https://adrenoleukodystrophy.info) lists 4163 variants, of which 90% pathogenic or likely pathogenic. These variants predominantly localize to the TM domain (46%) or the NBD (35%). The most frequent variant, p.Gln472Argfs*83, appears in no more than 5% of patients.^7^ It is estimated that 80-95% of cases are inherited, whereas the remainder result from *de novo* mutations.^8^ Data on approximately 10% of these variants indicate that at least 65% of missense variants lead to loss of function (LOF).^9^ Alongside roughly 18% frameshift or exon deletions, disease-causing mutations in *ABCD1* are classified as LOF variants, with no indications of disease-related gain-of-function (GOF) mutations.

ABCD1 is ubiquitously present in hepatocytes, kidney cells, skeletal muscle cells, endocrine systems, fibroblasts, macrophages, and endothelial cells. However, within the CNS, ABCD1 expression is confined to microglia, astrocytes, epithelial cells, certain oligodendrocyte subsets, and exclusively in neurons within the hypothalamus, basal nucleus of Meynert, periaqueductal gray matter, locus coeruleus, and dorsal root ganglia.^5,10,11^ Models for demyelination center on VLCFA-induced cytotoxicity amplified by the activity of VLCFA-specific elongase *ELOVL1* (converting C22:0 to C26:0),^1,12^ VLCFA’s role in proinflammatory macrophage activation,^13^ disturbance in calcium homeostasis triggered by VLCFA, and disrupted cholesterol homeostasis marked by increased cholesterol esters of saturated VLCFA and mono/polyunsaturated very- and long-chain fatty acids.^14^ For AMN axonopathy, models highlight oxidative stress and mitochondrial dysfunction,^15^ exacerbated by VLCFA cytotoxicity,^1^ and proinflammatory monocytes.^4^ Unlike peroxisomal biogenesis disorders, X-ALD does not appear to have a global impact on peroxisomal pathways and is thought to represent an isolated VLCFA defect. However, ABCD2, a peroxisomal ABC half-transporter highly homologous to ABCD1, may potentially alleviate peroxisomal pathology resulting from *ABCD1* LOF. It demonstrates shared substrate specificity, can heterodimerize with ABCD1, and its overexpression in mice is shown to reduce VLCFA accumulation.^5,16^ Nonetheless, while ABCD2 was found to modify metabolic impairment in mice,^17^ and it is not considered a phenotypic modifier in humans.^18^

Another challenge in X-ALD research is the difficulty of replicating X-ALD pathology in animal models. Despite ABCD1’s high interspecies conservation, animal models fail to mimic the inflammatory demyelination observed in ccALD.^5^ *Abcd1* hemizygous null mice show axonal degeneration of the sciatic nerve and spinal cord long tracts and hypermyelination. These symptoms have earlier onsets in *a Abcd1:Abcd2* double null mouse. *In vitro*, oligodendrocytes of Abcd1 null mice showed increased death rate during induced demyelination.^19^ Biochemically, elevated VLCFA levels are noted across various tissues, including a 2-8-fold increase in hexacosanoic acid,^5,20^ yet no diminution in peroxisomal β-oxidation activity was evidenced in mouse fibroblasts,^21^ as opposed to human fibroblasts.^22^ Additionally, these models do not exhibit adrenal pathology analogous to that seen in human X-ALD. Zebrafish models have demonstrated oligodendrocyte loss and myelin degradation with a modest VLCFA elevation (1.4-1.9-fold increase in C26:0) with unaltered cortisol levels. Nematode models deficient in *pmp-4* exhibit axonal degeneration affecting locomotion and a 25% increase in C26:0.

*Drosophila melanogaster* is one of the fundamental genetic models and has been extraordinarily successful in recent studies of rare and undiagnosed genetic disorders ^23,24^. Our group has also specifically studied peroxisomal genes in flies leading to novel disease insights and establishing *Drosophila* as a key pre-clinical model for peroxisomal disease ^25–28^.

Detailed studies of X-ALD pathogenic mechanisms in *Drosophila* could provide complementary insights to the existing vertebrate models ^5^. First, related peroxisomal genes such as ACOX1 have been studied in fly models, uncovering a novel human genetic disorder, Mitchell syndrome.^28^ Also, as previously noted, *Abcd2* may compensate in mice, but in flies there is a single *Drosophila* ortholog *Abcd1* (*CG2316*) for both vertebrate *ABCD1* and *ABCD2*. However, the *Drosophila* gene is on the 4th chromosome and has not been extensively studied. Preliminary work has shown that ubiquitous expression of Abcd1 RNAi induced defects in pigment glia (glial cells that support photoreceptors) and led to progressive loss of photoreceptors.^29^ Additionally, other genes in the fly, namely bubblegum (bgm) and double-bubble (dbb), a pair of homologous acyl-CoA synthetases that catalyze VLCFA esterification to CoA preceding Abcd1 transport activity, have been studied to gain insight into ALD. *bgm* homozygous null mutants displayed vision impairments linked to optic lobe anomalies, characterized by numerous vacuoles and the depletion of photoreceptors and pigment cells. These impairments were largely mitigated by dietary intervention with glyceryl trioleate oil, one of the two fatty acid components in Lorenzo’s oil that was originally investigated for the treatment of X-ALD and works by preventing the excessive accumulation of VLCFA. Furthermore, a double knockout of bgm and dbb resulted in significantly amplified neurodegeneration within the eye, manifesting as acute loss of lipid- and membrane-rich pigment photoreceptors, and adjacent glia ^30^.

In this study, we use knockdown and loss-of-function mutations in the fly *Abcd1* gene, and we designed transgenic flies that overexpress the human *ABCD1* gene. We report a range of phenotypes and explore the implications on peroxisome dynamics and function.

## MATERIALS AND METHODS

### Fly strains

Files were maintained in standard bottles containing water, yeast, soy flour, cornmeal, agar, molasses, and propionic acid as food at room temperature (22^0^C). Flies were raised either at room temperature, 25^0^C, or 29^0^C. Crosses of flies and 3^rd^ instar larvae collection were performed by standard procedures. The following *Drosophila* fly stocks were used:

*daughterless-da-GAL4*(P{GAL4-da.G32}UH1BL#95282, see below),

*eyeless-ey-GAL4* (BL#8228, y[1] w[1118]; P{w[+mC]=ey1x-GAL4.Exel}2),

*rh1-GAL4* (BL#8691, P{ry[+t7.2]=rh1-GAL4}3, ry[506]),

*elav*-*GAL4* (BL#8765, P{w[+mC]=GAL4-elav.L}2/CyO), and

*repo-GAL4* (BL#7415, w[1118]; P{w[+m*]=GAL4}repo/TM3, Sb[1]).

*UAS-luciferase RNAi* (BL# 35788, y[1] v[1]; P{y[+t7.7] v[+t1.8]=UAS-LUC.VALIUM10}attP2).

*UAS-lacZ RNAi* gift from Dr. Hugo Bellen lab

*UAS-eYFPPTS1* gift from Joseph Faust and James McNew at Rice University.

Two strains were used as control: Canton-S (gift by Dr. Hugo Bellen), and *y w* (BL#1495, y[1]w[1]).

*daGAL4, UAS-eYFP-PTS1* recombinant line made by Dr. Hyunglok Chung in Dr. Hugo Bellen’s lab.

dsRNA lines targeting dABCD1 were obtained from Vienna Drosophila Resource Center using the VALIUM20 vector,

*Abcd1RNAi^GD^ :(P{GD3151}v12170,*

*Abcd1RNAi^KK^ :(P{KK101414}VIE-260B)*.

### Generation of transgenic flies of humanABCD1 cDNA

Human *ABCD1-cDNA* clone IOH9909 from Ken Scott clone collection was utilized to generate transgenic flies. Mutations were generated in the same cDNA by using Q5 mutagenesis technique (NEB-E0554S) and confirmed by Sanger sequencing. Mutated plasmids were then injected in fly embryos to generate:

*y[1] w[*]; PBac{y[+mDint2] w[+mC]=UAS-hABCD1}VK00033/TM3,Sb[1]*

*y[1] w[*]; PBac{y[+mDint2] w[+mC]=UAS-ABCD1p.R518Q}VK00033/TM3,Sb[1]*

### Generation of *Abcd1* null(Primers selected *Abcd1^Δ4^*) flies

We generated a genomic loss-of-function variant by introducing a four-nucleotide deletion into the first exon of *Abcd1*. The line was made through the CRISPR pipeline for null mutants for the community by the NIG.

### Over-expression assay (Assessment of lethality and morphological phenotypes)

Lethality and morphological phenotyping assays were performed by crossing GAL4 driver (as indicated in the text) 5-10 virgin females to a similar number of males. After every 3-5 days, parents were transferred into a new vial to collect multiple F1 progenies. Flies were collected after most pupae were enclosed, and the total number of flies was scored based on the presence or absence of balancers. For the lethality assessment, a minimum of 100 flies were scored.

### Larval Assessment

For larval assay, 3^rd^ instar larvae F1 progenies were also isolated based on the lack of the Tubby or GFP balancer marker.

### RNA Isolation and Real Time Quantitative PCR

For each sample, 10 third-instar larvae were homogenized using the disposable Pellet Pestle (Sigma-Aldrich BAF199230001). RNA was extracted from the homogenate using 1ml TRIzol Reagent (Thermo Fisher, No. 15596018), followed by centrifugation at 12,000g for 12 minutes at 4°C (Eppendorf Centrifuge 5424R; Hamburg, Germany), retrieving the supernatant and discarding the pellet. The supernatant was incubated with chloroform followed by centrifugation (12,000g for 12 minutes at 4°C), and the aqueous phase was transferred to 100% isopropanol. Following incubation, samples were centrifuged (12,000 x g for 10 minutes at 4°C). The supernatant was discarded, and the tube was washed with 75% ethanol. Samples were centrifuged (7,500g for 5 minutes at 4°C) and air-dried for 20 minutes. RNA was quantified using the DeNovix DS-11 FX Nanodrop (DeNovix, Wilmington, DE), following the manufacturer’s instructions.

Real-time quantitative reverse transcription-PCR (real-time qRT-PCR) was performed to measure the mRNA levels of the studied genes using the CFX 96DX Touch real-time PCR detection system (Bio-Rad, Hercules, CA, USA). mRNA from 1 μg of total RNA was reverse transcribed into cDNA using the qScript™ cDNA Synthesis Kit Cat. No. 95047-025 (Quanta BioSciences, Beverly, MA), following the manufacturer’s instructions. qRT-PCR amplification was performed using the iTaq Universal SYBR^®^ Green (Bio-Rad, Hercules, CA, USA), following the recommendations of the manufacturer. For expression studies, the qRT-PCR results were normalized against an internal control (RPL32). The primers were designed using the Primer 3 Plus software^31^ with the following parameters: product size of 70 to 150 base pairs, melting temperature (Tm) of 58 to 62°C, length of 18 to 23 nucleotides, and GC content of 30 to 80%.

Primers selected:

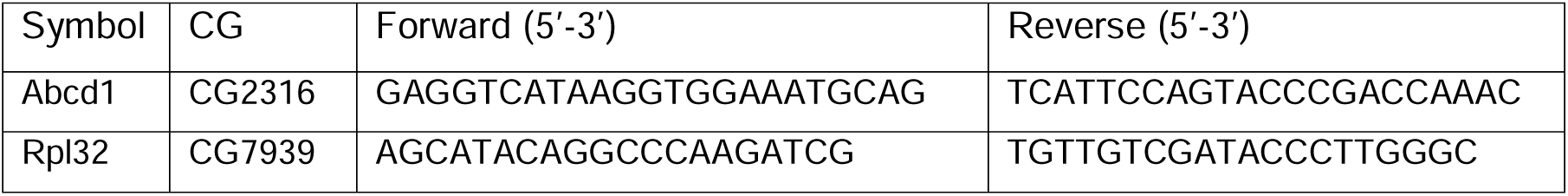

### Locomotion assays

A robotic system facilitating the negative geotaxis climbing assay was utilized as previously described.^32^ In brief, a custom robotic geotaxis apparatus was designed at the Automated Behavioral Core of the Dan and Jan Duncan Neurological Research Institute (led by Dr. Juan Botas) to convert the locomotor behavior of flies into measurable data. This apparatus taps Drosophila encased in arrays of 96 vials, with their movements within these vials captured at a rate of 30 frames per second. Each genotype was represented by 5 replicates, and each vial contained 8-11 adult flies, observed at intervals of 3-4 days. The metrics recorded included the flies’ distance traveled, velocity (mm/sec), turning, and the number of stumbles within the vials. Each genotype contained at least 30 flies.

### Lifespan assays

Lifespan analysis was performed as previously described.^28^ Briefly, newly eclosed flies were separated by genotype and sex and incubated at 25^0^C. Flies were transferred to a fresh vial every two days and survival was determined once a day. 50 flies were tested per group. Statistical analysis was performed using the Log-rank (Mantel-Cox) test.

### Nile red staining

For lipid droplet staining, fly heads were dissected in cold (4°C) phosphate-buffered saline (PBS) and fixed in 4% paraformaldehyde (Merck, Darmstadt, Germany) in PBS for 20 minutes. Retinas were then dissected under PBS and rinsed three times in 1X PBS on a rotating platform, for 10 minutes in each wash. Samples were then incubated for 30 min in 1 mg/mL Nile red (Sigma, N3013) (1:1000 dilution with PBS) and then rinsed three times with 1X PBS. Stained samples were mounted on a glass slide in VectaShield (Vector Laboratories, #H1000) for imaging on a Leica SP8 confocal microscope. Images were obtained using a 40x and/or 63x glycerol immersion lens with 3x zoom.

### Electroretinograms

Electroretinogram (ERG) recordings were performed as previously described.^33^ In brief, female flies were glued to a slide with Elmer’s Glue. A recording electrode filled with 100 mM NaCl was placed on the eye, and a reference electrode was placed on the fly head. During the recording, a 1 s pulse of light stimulation was given, and the ERG traces of eight flies for each genotype were analyzed with pCLAMP 9.2 software (Molecular Devices). Values for amplitudes and depolarization amplitudes (’on and off’ transients) (in mV) represent an average of a total of at least 24 electrograms per genotype.

### Immunocytochemistry

Staining of peroxisomes in the salivary gland was performed as previously described.^34^ Briefly, fresh third-instar larvae were dissected, and salivary glands were visually identified. The salivary glands were separated and fixed with 4% paraformaldehyde in PBS for 15 minutes. They were then permeabilized with 1% Triton X-100 in PBS (PBTx) at room temperature by washing three times for 10 minutes each on a rotating platform. Subsequently, samples were incubated overnight at 4°C on a rotating platform with Rabbit-anti-Pex3 antibody (1:500 ^27^). Then the samples were washed again with 1% PBTx three times for 10 mins each. These samples were then treated with anti-rabbit Alexa Fluorophore Cy3 (Invitrogen #A-11011, 1:500), and with 4′,6-diamidino-2-phenylindole (DAPI, 1:1000) secondary antibodies diluted in 1% PBTx for 2 hrs. at room temperature. Following incubation, the samples were washed four additional times, 10 minutes each, with 1% PBTx. Then the samples were mounted on a glass slide with VectaShield (Vector Laboratories, #H-1000) for imaging on a Zeiss 710 confocal microscope. Images were obtained using a 63x - oil immersion lens with 1.5x digital magnification.

### Quantitative Analysis of Fluorescence Images

All fluorescence images were quantified and analyzed using the Surface module of Imaris v9.8.2 (Bitplane, Zurich, Switzerland). In this module, a surface was generated for the fluorophore signal of interest by applying background subtraction and setting a lower area filter limit of 0.5 µm². This threshold was chosen to reduce background noise and prevent false positives during surface creation. The number of puncta (surfaces created) in a single cell was determined from the statistical data provided for the surfaces.

### Peroxisomal Biochemical Assays

VLCFA analysis was performed on third-instar larvae, separated by genotype. Ten larvae of each phenotype were collected and placed in 1.5 mL Eppendorf safe lock microtubes. The larvae were rapidly rinsed in 1X PBS three times to remove food residues. The tubes were then flash-frozen in liquid nitrogen for 5 seconds and stored in a standard −80°C freezer. Very long chain fatty acid quantification was performed with gas chromatography and liquid chromatography mass spectrometry (GC-MS and LC-MSMS) as previously described (Dr. Ann Moser, Kennedy Krieger Institute).^35^ In brief, a modified Folch extraction was used to obtain lipids. Lipid fatty acids were acid hydrolyzed from triglycerides and phospholipids then measured as their pentafluorobenzyl bromide esters by isotope dilution capillary gas chromatography negative chemical ion GCMS (see Lagerstedt et. al. 2001^36^). Lysophosphatidylcholine were analyzed by LC-MSMS on an AB Sciex API3200 or QTRAP 4500 with a Phenomenex Kinetex C8 column (2.6μm x 50 x 2.1 mm) using a linear gradient between 0.1% formic acid in H_2_O and 0.1% formic acid in methanol in positive electrospray ionization mode (see van de Beek et. al.^37^) For each genotype, prior to homogenization, five larvae were allocated for GC-MS and four for LC-MS, with 4-6 replicate tubes per genotype (N=4 to 6).

### Statistical analysis

For the ERG experiments, a 2-tailed Student’s t-test was used to assess the statistical significance of differences between analyzed groups. Results are presented as means ± standard deviation (SD). For locomotion analysis, individual timepoints from the three datasets were directly compared using multiple unpaired parametric Welch t-test analysis with a two-stage Benjamini, Krieger and Yekuteili step-up for false discovery rate correction. For lipidomic analysis, multiple comparisons were performed using the Brown-Forsythe and Welch ANOVA tests, followed by a post hoc Dunnett’s T3 multiple comparison test between the different genotypes and a *y^1^ w\** control. Throughout the text, a p-value < 0.05 was considered significant and marked by *, a p value < 0.01 by **, and a p value < 0.001 by ***, and a p value < 0.0001 by ****; for the locomotion analyses (figure 2) either *, #, or ** marked a p-value ≤ 0.0001 depending on the compared dataset. Higher p values were not marked (see figure caption).

## RESULTS

### Abcd1 knockdown resulted in salivary gland defects, retinal lipid abnormalities and reduced survival

*Drosophila Abcd1* (*CG2316*, Flybase ID: FBgn0039890) is a 730-amino-acid polypeptide that shows 71% similarity, 52% identity, and 4% gap to human *ABCD1*^38^ (**Supplementary Figure 1A-C**). To assess the loss of function of *Abcd1*, the fly’s only CoA-esterified long-chain fatty acids peroxisomal importer, we tested three RNAi stocks, but we pursued only two as the third one didn’t show the knockdown effect (**Supplementary Figure 1D**). The *UAS-RNAi* were ubiquitously expressed using the GAL4/UAS system with the transcriptional activator GAL4 linked to a *daughterless (da)* driver, an early and broadly expressed gene essential for proper embryonic development,^39^ to drive the expression of dsRNA for *Abcd1* attached to a UAS enhancer. For the optimization of the GAL4/UAS expression system, flies were maintained at 29^0^C with a 12-hour light/dark cycle. These flies also expressed the peroxisomal matrix marker (*eYFP-PTS1*).^26,27^. Both *UAS-Abcd1-RNAi* being on X-chromosome we used males to cross with *da-GAL4-UAS-eYFPPTS1* females for the knockout assay. We observed that overexpression of *Abcd1^GD31^*^51^ (henceforth *Abcd1-RNAi^GD^*) showed a 90% decrease in *Abcd* mRNA expression, while *Abcd1^KK1014^*^14^ (henceforth *Abcd1^-^RNAi^KK^*) demonstrated a 50% reduction (**Figure 1A**). Both RNAi strains were predicted to have no off-target effects.

**Figure1.**
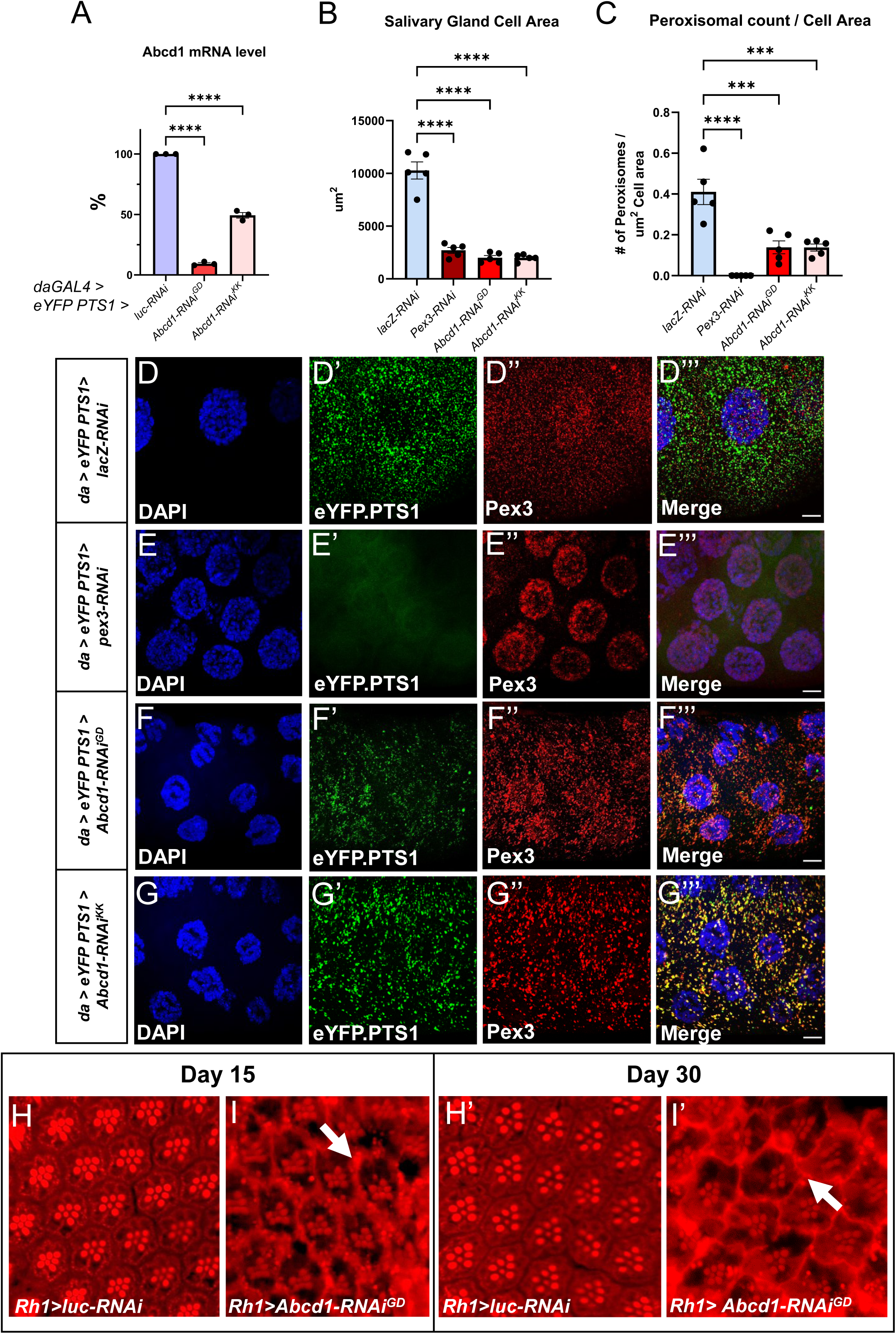
*Drosophila Abcd1* knockdown affects cell size and peroxisome abundance in Salivary gland. *da-GAL4* is used to overexpress both the peroxisomal marker *eYFP.PTS1* and the appropriate RNAi lines ubiquitously at 29^0^C. (**A**) Total Abcd1-mRNA reduction in larvae overexpressing *Abcd1-RNAi* GD3151 (*Abcd1-RNAi^GD^*) and KK101414 (*Abcd1-RNAi^KK^*) showed different levels of mRNA knockdown as compared to *Luc-RNAi* control. *Abcd1-RNAi^GD^* displayed a greater degree of knockdown than *Abcd1-RNAi^KK^*. (**B**) Ubiquitous *Abcd1* knockdown via *da-GAL4* decreases the size of third instar salivary glands compared to *lacZ-RNAi* control. It is very similar to that is observed in *pex3*-knockdown salivary glands. (**C**) Utilization of peroxisomal membrane marker *eYFP.PTS1* to visualize the peroxisomes in the salivary gland showed significant reduction in peroxisomal abundance in both *Abcd1-RNAi* knockdown fly strains. Imaris software is utilized to calculate the number of peroxisomes per cell area (um^2^). (**D-G’’’**) Third instar larval salivary gland images of peroxisomes along with *a*nti-Pex3 antibody staining. *daGAL4>UAS-eYFP.PTS1* flies expressing *UAS-lacZ-RNAi* in panel **D-D’’’**, *UAS-pex3-RNAi* in panel **E-E’’’**, *UAS*-*Abcd1-RNAi^GD^* in panel **F-F’’’**, and UAS-*Abcd1-RNAi^KK^* in panel **G-G’’’**. These panels show the separate single channel confocal images of nucleus stained with DAPI, peroxisomes marked with *eYFP.PTS1* and Pex3 protein stained with Cy3 along with the merge in the right most panel with 10um scale bar. ImageJ software was utilized to generate the maximum intensity projection and the scale bar in Merge panels. (**H-I’**) Retinal cross section of adult eyes stained with Red Nile at the age of 15 days after eclosion (DAE) (lefts panels) and 30 DAE (right panels). The *Rh1-GAL4 > UAS-Luc-RNAi* control flies are shown in **H** and **H’** and *Rh1>Abcd1-RNAi^GD^*flies are shown in **I** and **I’**. The lipid droplets within the ommatidial structure are shown with the white arrows. [*ns* – nonsignificant, indicates p value ≥ 0.05; ** indicates a p-value < 0.01, **** indicates a p-value < 0.0001.]

Reduction of *Abcd1* translation was associated with significant reduction in viability (**Supplementary Table 1**). Crosses with *Abcd1-RNAi^GD^* resulted in female adult lethality, yielding only a few survivals. Moreover, the female larvae reached third instar stage but exhibit a prolonged and terminal larval stage, did not pupariate on time (**Supplementary Figure 2**), and they are lethal in early pupal stage with very few survivors. This result indicates that reduction *Abcd1* RNA levels affect survival. Furthermore, surviving adult flies for *Abcd1-RNAi^KK^* had reduced lifespan and female fertility (**Supplementary Table 1**) compared with control.

We next moved to characterize phenotypic abnormalities in larvae and adult flies (**Figure 1**). For larval analysis, we again used *da-GAL4-UAS-eYFPPTS1* to drive *Abcd1-RNAi^GD^*and *Abcd1-RNAi^KK^*, and we utilized *lacZ-RNAi* as a negative control. We also compared these with *pex3-RNAi* which we previously established as disruptive of peroxisomal biogenesis.^27^ Flies reared at 29°C and we dissected *Drosophila* third instar larvae salivary glands from these F1 female larval progenies (**Figure 1B-C Quantification of images Figures 1D-G**) and observed that the salivary glands in *pex3* knockdown larvae were smaller due to reduced cell size; this effect was similarly observed in the *Abcd1* knockdown (**Figure 1B**). We also expressed a peroxisomal matrix marker (*eYFP-PTS1*) that targets the peroxisomes and used anti-Pex3 antibody to immunofluorescently stain the membrane of early and mature peroxisomes.^26,27^ As expected, *pex3* knockdown led to total loss of peroxisomal localization of the YFP-PTS1 and presumed non-specific nuclear staining with the anti-Pex3 antibody (**Figure 1E**). Knockdown of *Abcd1* with either of the RNAi transgenes did not disrupt the peroxisomal punctae where YFP-PTS1 and Pex3 protein co-localize (**Figure 1F-G**). Additionally, there is a significant reduction in the peroxisomal abundance in the larvae’s salivary glands compared to controls (**Figure 1C**). In summary, ubiquitous knockdown of *Abcd1* led to poorly developed salivary glands with small cells similar to knockdown of *pex3*. Knockdown of *Abcd1* did not eliminate peroxisomes as *pex3* knockdown but did appear to reduce the number of peroxisomes per cell area.

For adult analysis, we expressed *Abcd1-RNAi^GD^* it with a *Rh1-GAL4* driver, a G protein-coupled receptor encoded by ninaE and expressed in the R1-R6 photoreceptor cells in the compound eye. Flies were reared at 29°C This resulted in viable adults but with an abnormal appearance of the ommatidia and abnormal distribution of Nile staining indicating an alteration in lipid droplets at 15 days after eclosion (written henceforth as DAE) compared to control (**Figure 1H-I**). Moreover, by 30 DAE, this phenotype appears to progress and worsen: the retina had more marked lipid droplet alterations (**Figure H’-I’**). Electroretinogram (ERG) of *Rh1>Abcd1-RNAi^GD^*on 35 DAE showed a 17% increased depolarization amplitude, and a 14% and 37% reduction in the on- and off-transients by respectively, when compared to *Rh1>lacZ-RNAi* (**Supplementary Figure 3**). No significant ERG changes were observed for *Eyeless-GAL4 driven Abcd1-RNAi^GD^* line, other neuronal GAL4 drivers which included the pan-glial repo, and the pan-neuronal elav (data not shown).

### Genomic out-of-frame deletion of Abcd1 resulted in age-related locomotion impairment

We also generated a genomic loss-of-function variant of *Abcd1* by introducing a four-nucleotide deletion into its first exon in *y^1^, w\** flies, named as *Abcd1*^Δ*4*^ (written as SKΔ4 in **Supplementary Figure 1D**). The deletion was verified by amplification failure (**Supplementary Figure 4**). This *Abcd1*^Δ*4*^ genomic variant, did not demonstrate reduced fertility and showed no statistically significant reduction in survival compared to *y^1^,w\**(henceforth written as *yw*) flies (data not shown). Similarly, we did not observe gross salivary gland pathology. To assess the motor effects on the adult flies, we employed a specialized robotic system designed to capture video footage of flies as they ascend to the peak of a vial after being displaced to the base. The movement of individual flies is monitored in real-time, and we assessed several locomotor metrics. *Abcd1*^Δ*4*^ homozygotes (*Abcd1*^Δ*4/*Δ*4*^) were compared to *yw* and *Abcd1*^Δ*4*^ heterozygotes-*Abcd1*^Δ*4/+*^ (*Abcd1*^Δ*4*^ homozygotes crossed to *yw*). Relative to both *Abcd1*^Δ*4/+*^ and *yw* flies, *Abcd1*^Δ*4/*Δ*4*^ females exhibited a significant decrease in velocity that exacerbated with age (**figure 2A**). Specifically, a 21-31% reduction in velocity was observed within the initial 20 DAE, increasing to 33-49% by 30 DAE, and further intensifying to a 79% reduction by 43 DAE. Over the span from 7 to 43 DAE, these mutant homozygous flies demonstrated an 80% decrease in velocity, in contrast to a 45% reduction observed in heterozygotes, and a 35% reduction in *yw* flies. Additionally, *Abcd1*^Δ*4/*Δ*4*^ females displayed an age-dependent increase in stumbling events. No notable differences were observed among the three genotypes within the initial 22 DAE. However, from 26 to 43 DAE, *Abcd1*^Δ*4/*Δ*4*^ females experienced a 275% increase in stumbling events, while *Abcd1*^Δ*4/+*^ and *yw* genotypes only exhibited 152% and 145%, respectively (**figure 2B**). In comparison, *Abcd1*^Δ*4/*Δ*4*^ males also exhibited significant locomotor impairments, albeit less pronounced than in females, with velocity reductions ranging from 15% to 44% relative to controls. Over the span from 7 to 43 DAE, these mutant males experienced a 60% decrease in velocity, versus a 51-52% decrease observed in controls (**figure 2C**). Moreover, starting from 26 DAE, *Abcd1*^Δ*4/*Δ*4*^ males showed a 215% increase in stumbling events, compared to 147% in *Abcd1*^Δ*4/+*^ and 173% in *yw* genotypes (**figure 2D**). In summary, *Abcd1*^Δ*4/*Δ*4*^ did not show reduced survivability but exhibited age-dependent impaired locomotion, with more significant impairment in females than males.

**Figure 2.**
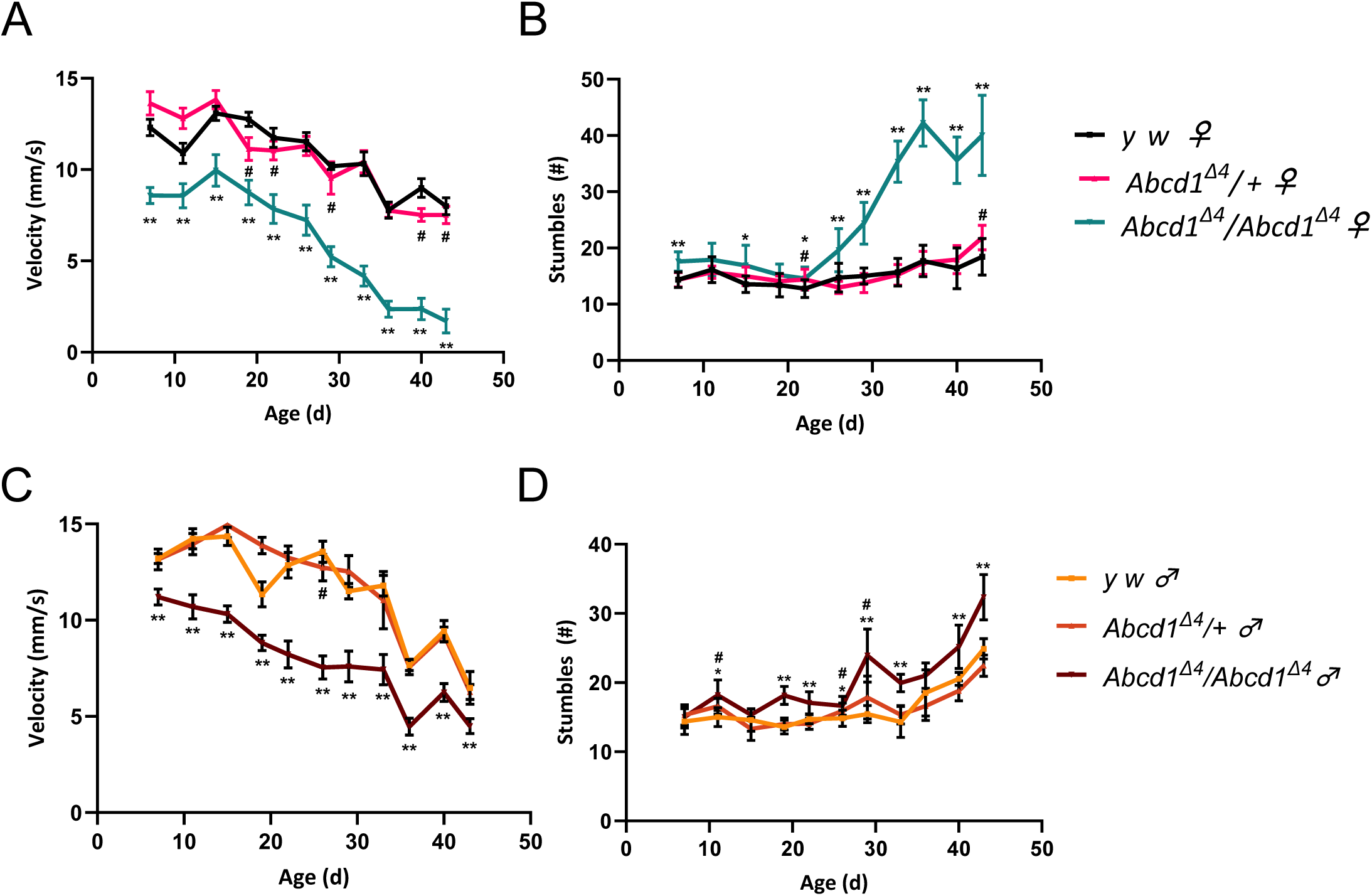
Locomotion impairment with out-of-frame deletion in the *Abcd1* gene. A robotic negative geotaxis system recorded climbing speed and stumbling in flies every 3-4 days starting at 7 DAE (see Materials and Methods). Results are separated by sex (left panels for females, right panels for males), N ≥ 25 for each sex and genotype. (**A**) The velocity of *Abcd1*^Δ*4/*Δ*4*^ (homozygous) females with the deletion is significantly reduced compared to *yw* controls and *Abcd1*^Δ*4/+*^ (heterozygotes), in an age-dependent manner. Reduction in younger females (first 20 days after eclosion DAE) amounted to 21% (19 DAE) and 31% (7 DAE), escalating to 49% at 29 DAE, and 79% at 43 DAE. (**B**) Stumbling events increased from a baseline similar to controls and heterozygotes up to 22 DAE, then increased by 2.4-2.9-fold in 36-43 DAE compared to an increase of 1.2-1.5-fold in age-matched controls and heterozygotes. (**C**) The velocity of homozygous males was similarly reduced compared to heterozygotes and controls (by 15-44%); however, the gap does not widen with age. (**D**) An age-dependent increase in stumbling is seen by up to 1.9-fold (from 22 DAE), more than heterozygotes and controls (in comparison to 1.6 and 1.7-fold increases, respectively). In every panel, across all timepoints, data were deemed statistically significant at a p-value ≤ 0.0001 (refer to Materials and Methods for calculation details). Homozygotes that performed significantly worse than controls are indicated by *, and ** when their performance is compared to both controls and heterozygotes. The # symbol denotes statistical significance of heterozygotes relative to controls.

### Abcd1 knockdown or loss-of-function caused the accumulation of saturated VLCFA in larvae

The biochemical hallmark in patients with X-linked adrenoleukodystrophy (X-ALD) is the accumulation of VLCFAs (C>22), particularly C26:0 (hexacosanoic acid), and to a lesser extent, C24:0 (tetracosanoic acid) and with nominal difference in C22:0 (docosanoic acid, not considered a VLCFA).^40^ Thus, we assessed for a similar biochemical signature in both the RNAi-mediated knockdown (*Abcd1-RNAi^GD^*and *Abcd1-RNAi^KK^*) with *da-GAL4* and the *Abcd1* out-of-frame small deletion (*Abcd1*^Δ*4/*Δ*4*^) larvae using gas chromatography-mass spectrometry (GCMS) (**Figure 3**). *Abcd1*^Δ*4/*Δ*4*^ was compared to both *yw* and heterozygotes (*Abcd1*^Δ*4/+*^), and *Abcd-RNAi* was compared with *yw* and *da>Luc* (*da>luciferase-RNAi*).

**Figure 3.**
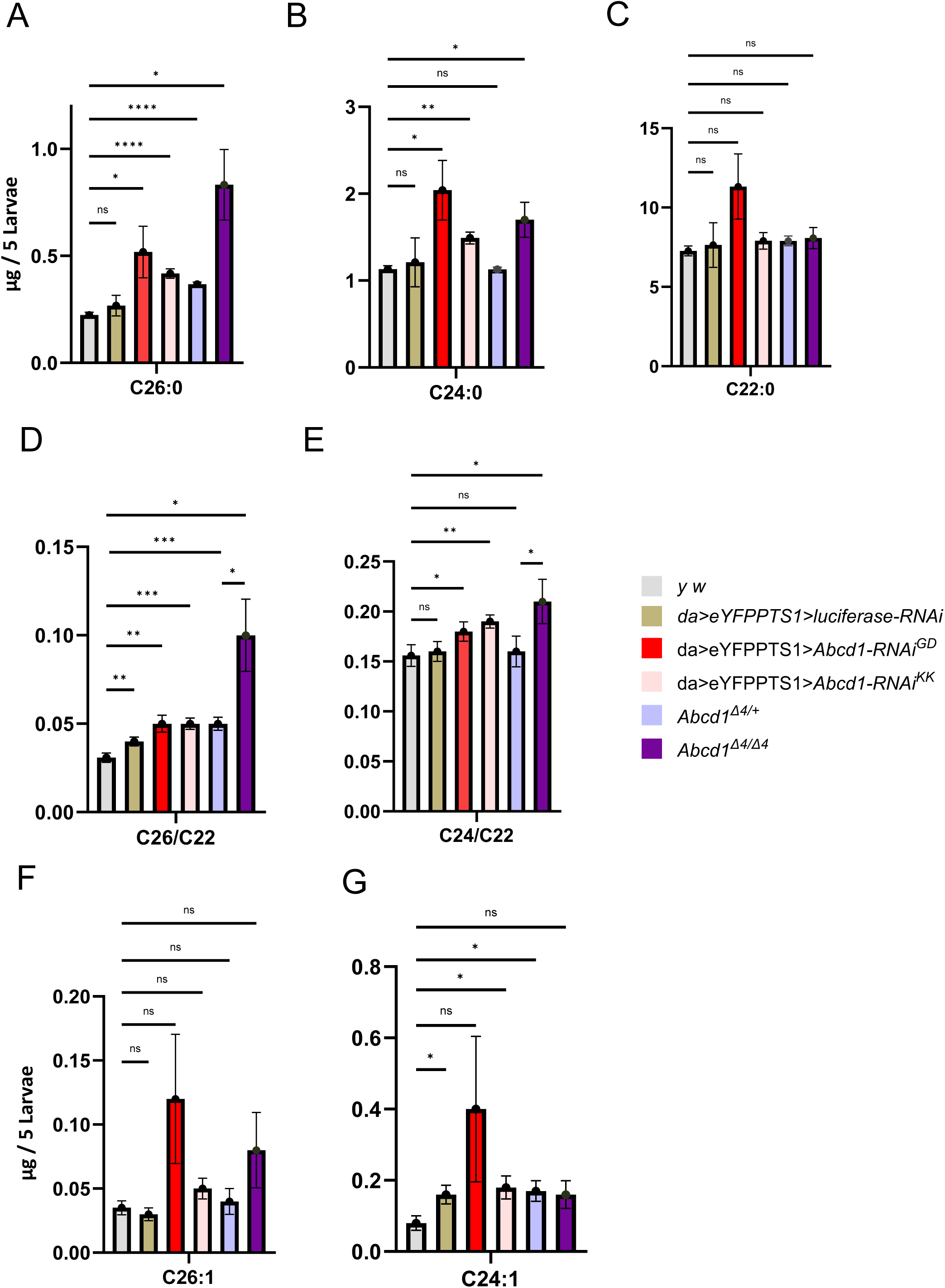
Accumulation of VLFCA in third instar larvae with *Abcd1* LOF. The quantification of hexacosanoic acid (C26:0) **(A)**, tetracosanoic acid (C24:0) **(B),** and docosanoic acid (C22:0) **(C)** in micrograms per 5 larvae. C26:0, a key marker for X-ALD, shows an increase in our deficiency models. *da>Abcd1-RNAi^GD^* accumulates C26:0 levels by 2.32- and 1.94-fold compared to *yw* and *da>luciferase^RNAi^* controls, and *da>Abcd1-RNAi^KK^*larvae result in a 1.9 and 1.6-fold increase, respectively. *Abcd1^Δ4/Δ4^* leads to a 3.7-fold increase in C26:0 levels compared to the *yw* control, and a 2.7-fold increase compared to deletion heterozygotes (*Abcd1*^Δ*4/+*^). Increases are also observed for C24:0, whereas significant elevations of C22:0 are only seen with *Abcd1-RNAi^GD^* compared to controls as shown in **(C),** however C22:0 elevations did not reach statistical significance. In **(D),** *Abcd1*^*Δ4/Δ4*^ showed increases in the ratios of C26:0/C22:0 by factors of 3.3 over controls, and 2.1 over heterozygotes. The RNAi knockdown resulted in milder, yet statistically significant, elevations in these ratios (1.3-1.7-fold for C26:0/C22:0 for both strains). **(E)** a similar trend is illustrated in the C24:0/C22:0 ratio. **(F-G)** illustrate the accumulation of C26:1 and C24:1 peroxisomal markers: C26:1 accumulated 3.5-3.7-fold more in *Abcd1-RNAi^GD^*compared to controls and a milder 1.4-1.5-fold increase in *Abcd1-RNAi^KK^* and 2.3-fold more in *Abcd1*^*Δ4/Δ4*^ compared to controls and heterozygotes. C24:1 accumulated in *Abcd1-RNAi^GD^* by a factor of 2.6-5 and in Abcd1-RNAi^KK^ by a factor of 1.2-2.2. However, these elevations of the monounsaturated VLCFA did not reach statistical significance. Lipids were isolated using GCMS. *ns* – nonsignificant, indicates p value ≥ 0.05; ** indicates a p-value < 0.01, **** indicates a p-value < 0.0001.

For RNAi, *Abcd1-RNAi^GD^*larvae exhibited an accumulation of total saturated VLCFA, particularly C26:0, which increased by 1.9-2.3-fold compared to controls (yw and da>Luc). The less potent RNAi strain, *Abcd1-RNAi^KK^*, showed a more moderate elevation of 1.6-1.9-fold, suggesting a dose-response relationship (**Figure 3A**). More significantly, *Abcd1*^*Δ4/Δ4*^ showed accumulation of C26:0: 3.7-fold compared to *yw*, and 2.7-fold compared to *Abcd1*^Δ*4/+*^ (**Figure 3A**), although the latter did not reach statistical significance (*p* value = 0.067). The *Abcd1-RNAi^GD^* strain significantly elevated C24:0 by 1.7-1.8-fold and C22:0 levels by 1.5-1.6-fold although the latter did not reach statistical significance. *Abcd1-RNAi^KK^* elevated only C24:0 levels, by 1.2-1.3-fold (**Figure 3B-C**). *Abcd1*^*Δ4/Δ4*^ also showed significant increases of 1.5-fold (compared both to yw and *Abcd1*^Δ*4/+*^) with no increase for C22:0 (**Figure 3B-C**).

Additionally, *Abcd1*^*Δ4/Δ4*^ showed increases in the ratios of C26:0/C22:0 and C24:0/C22:0 (commonly used in humans for diagnostic purposes^41^) by 3.3 fold and 1.4 fold over controls (*yw*), and 2.1 and 1.3 over heterozygotes (*Abcd1*^Δ*4/+*^), respectively. The RNAi knockdown resulted in milder, yet statistically significant, elevations in these ratios: 1.3-1.5-fold for *Abcd1-RNAi^GD^* for the C26:0/C22:0 ratio, and 1.5-1.7-fold for *Abcd1-RNAi^KK^*; and 1.15-1.21-fold increase for C24:0/C22:0 for both strains (**Figure 3D and 3E**).

C26:1 and C24:1, another VLCFA markers for peroxisomal function, showed trend towards accumulation in *Abcd1-RNAi^GD^* (3.5-3.7-fold fold for C26:1 and 2.6-5-fold for C24:1), in *Abcd1-RNAi^KK^* (1.4-1.5-fold and 1.2-2.2-fold, respectively) and *Abcd1*^Δ^*^4/^*^Δ^*^4^* (2.3-fold and 1-1.9-fold) compared to controls, although these changes did not reach statistical significance (**Figure 3F and G**).

Overall, these results indicate that VLCFA is elevated in both knockdown (*Abcd1-RNAi^GD^, Abcd1-RNAi^KK^)* and genomic loss-of-function variant in Abcd1 (*Abcd1*^Δ^*^4/^*^Δ^*^4^*), with a dose-response effect in the RNAi strains.

Replication of these findings by LCMSMS (**Supplementary Figure 5**) revealed even more pronounced elevations in C26:0-carnitine with 3.3-3.5-fold and 3.4-3.6-fold increases for both RNAi strains, and a 3.2-fold increase in *Abcd1*^*Δ4/Δ4*^ with only 1.3-fold elevation in *Abcd1*^Δ*4/+*^. Levels of C26:0, C24:0, and C22:0 (grouped) were also elevated, by factors of 1.7-3.5 in the RNAi, and 3.6 in *Abcd1*^*Δ4/Δ4*^ (**Supplementary Figure 5**). Results indicated a trend towards mildly increased levels of LPCs, which did not reach statistical significance (**Supplementary** Fig. 5). Overall, our model aligns with the primary biochemical finding of hexacosanoic acid elevation in *Abcd1* loss-of-function, as observed in human disease.

### Overexpression of human ABCD1 gene caused a semi-lethal phenotype with abnormal retinal ultrastructure, and wing morphology

Having observed loss-of-function phenotypes in *Drosophila Abcd1* deletion and knockdown, we designed a transgenic human *UAS-ABDC1* to attempt rescue. In parallel, we selected a disease-causing *ABDC1* missense mutation, a recurrent pathogenic variant p.Arg518Gln, located in the highly conserved fifth exon (**Supplementary Figure 6**) (NM_000033.4(ABCD1):c.1553G>A (p.Arg518Gln), ClinVar accession: RCV000077955) (**Supplementary Table 1**). However, ultimately, we were unable to perform rescue as we observed toxicity from the expression of the human transgenes.

Because we had observed LOF retinal phenotypes, we first evaluated overexpression in retina (in flies with endogenous CG2316 intact) and expressed both human UAS-*ABCD1^Ref^* and UAS-*ABCD1^R518Q^*using the *Rh1-GAL4* driver (*Rh1*). Flies were reared at 29^0^C. The retina for these flies showed markedly abnormal architecture with vacuoles and ommatidial disarray at 15 DAE compared to control (*lacZ-RNAi*) (**Figure 4B compared to 4A)**. Interestingly, the *ABCD1^R518Q^* loss-of-function variant did not appear to affect retinal structure (**Figure 4C**). We also performed ERG at 15 DAE and observed a statistically significant reduction in amplitude, and depolarizing on and off transients, by 21%, 48%, and 54% respectively (**Figure 4D and Supplementary Figure 3**).

**Figure 4.**
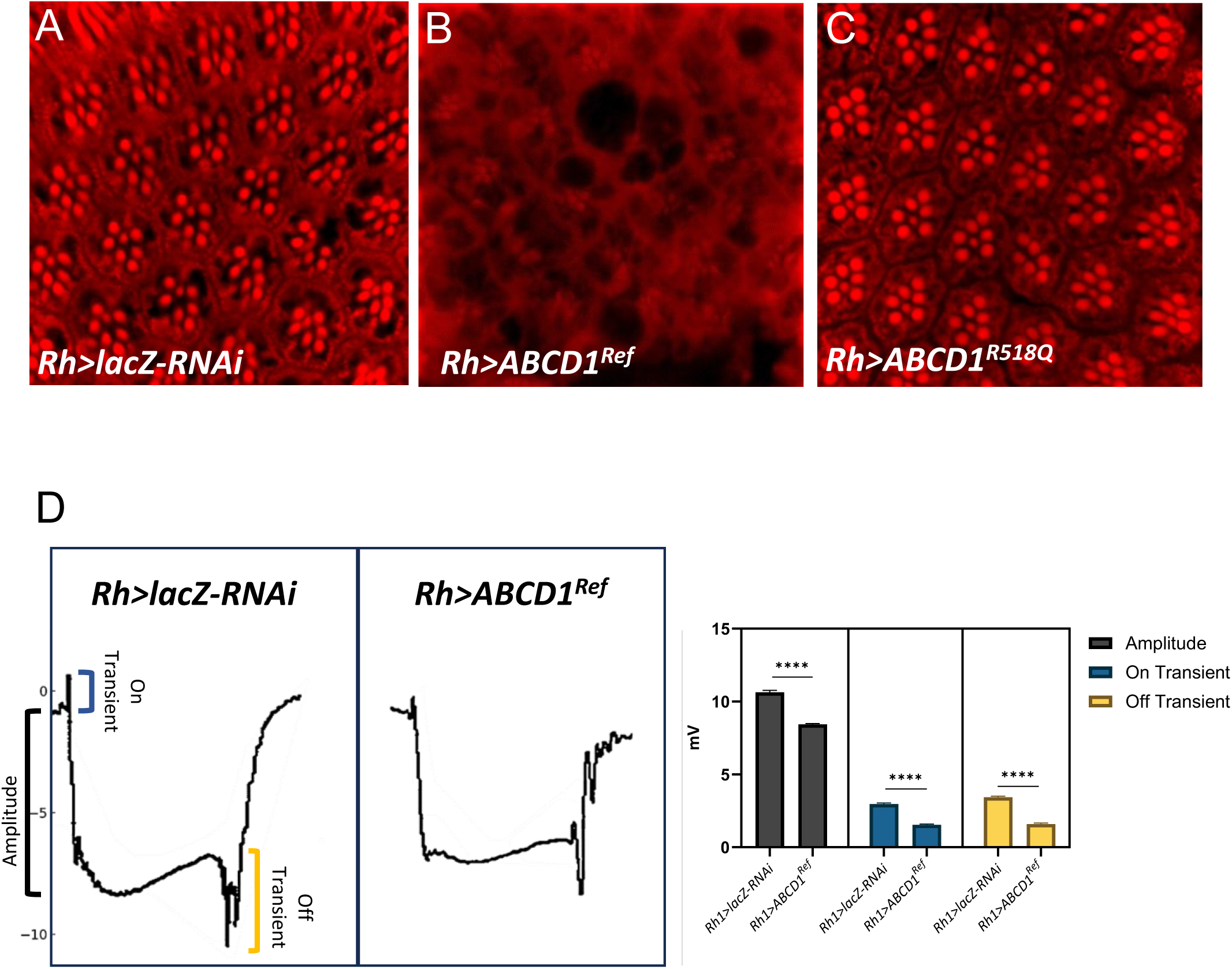
Overexpression of Human ABCD1 caused abnormal retinal morphology. **(A-C)** Human ABCD1 transgenes are overexpressed utilizing *Rh1-GAL4* driver and adult flies are imaged at 15 days after eclosion (DAE). Overexpression of human *ABCD1^Ref^* in panel **(B)** showed very disorganized lipid morphology relative to the control *lacZ-RNAi* **(A)**. However, *ABCD1^R518Q^* did not display defects in retinal morphology as shown in panel **(C)**. **(D)** Electroretinogram assay performed at 15 DAE on the flies expressing human *Rh1>ABCD1^Ref^*showed a significant reduction in Amplitude, On transient and Off transient as compared to *Rh1>lacZ*.

We then overexpressed the human proteins *ABCD1^Ref^* and *ABCD1^R518Q^* using *daughterless* driver (*da-GAL4, UAS-eYFP.PTS1*). Flies were also reared at 29^0^C. We observed some reduction in the survival of flies overexpressing *ABCD1^Ref^* in comparison to *lacZ-RNAi* with the same driver (*da>lacZ-RNAi* controls) (**Supplementary Table 1, Supplementary Figure 2**). Adult *ABCD1^Ref^*female flies showed reduced fertility, with only 3 progenies per female. Interestingly, about 50% of the flies showed a crumbled wing phenotype compared to control flies (**Figure 5C compared to Figure 5A**). In addition, none of them survived more than 20 days after eclosion (DAE). Flies overexpressing the p.R518Q missense pathogenic variant *ABCD1*^R518Q^, also reared at 29^0^C, demonstrated increased survival compared to *ABCD1^Ref^*with 2 viable offspring per female per 24 hours, reflecting a 2.5-fold improvement compared to *ABCD1^Ref^*. Female fertility improved similarly by 2.7-fold to 8 viable offspring for each female (**Supplementary Table 1**). 25% of the flies survived to 30 DAE, reaching up to 41 days, in comparison to 0% for *ABCD1^Ref^*. Overexpression of the *ABCD1^R518Q^* variant did not produce wing crumpling. Wing crumpling is a phenotype that we have observed in *Drosophila pex* gene mutations and indeed we have observed this phenotype with *pex2* mutants (**Figure 5B**). This data suggests that overexpression of the *ABCD1* human reference (but not p.R518Q mutant) protein produces phenotypes related to defects in peroxisomal biogenesis defects in flies.

**Figure 5.**
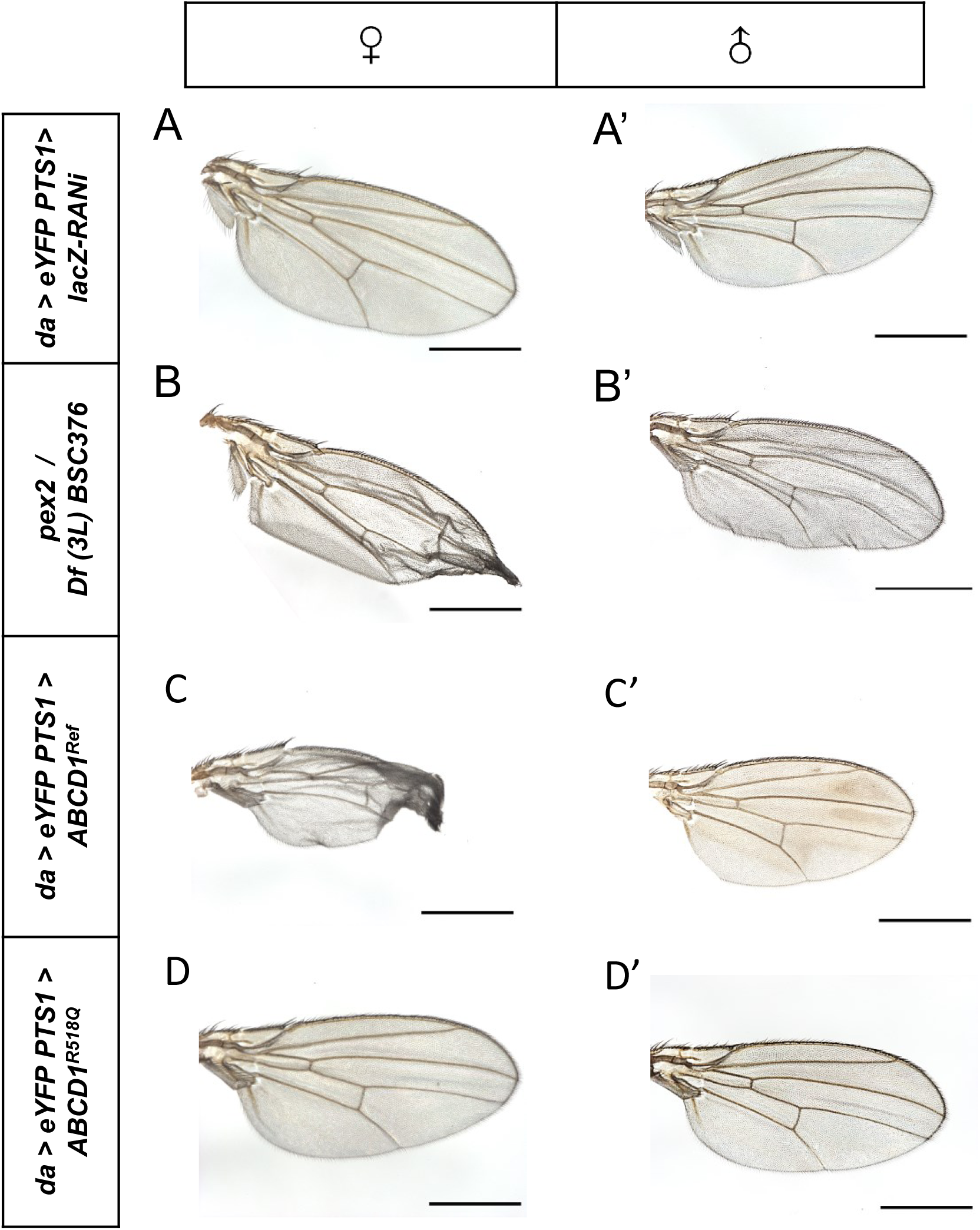
Crumpled-wings observed in ubiquitous overexpression of adults expressing human *ABCD1^Ref^*mimicked the crumpled-wings seen in *pex2* LOF. Ubiquitous overexpression of *ABCD1^Ref^* demonstrated crumbled-wing phenotype compared to control flies **(C) compared to (A)**. Similar wing phenotype is seen in *pex2* loss of function (**B**). The variant *ABCD1^R518Q^* expressing flies in panel (**D**) did not display the same wing phenotype.

In order to test this, we imaged peroxisomes in third instar larvae salivary glands. Flies were reared at 29^0^C. Again, we observed loss of eYFP-PTS1 localization and non-specific Anti-Pex3 antibody staining in the *pex3-RNAi* knockdown compared to controls (**Figure 6B compared to A**). Larvae in which human *ABCD1^Ref^* is overexpressed revealed grossly normal appearing salivary glands although there was a small difference in cell size (**Supplementary Figure 7**). However, peroxisomal staining with YFP-PTS1 matrix marker and Anti-Pex3 antibody revealed a striking loss of peroxisomes (**Figure 6C**). While there was some similarity to the *pex3-RNAi* knockdown we did observe occasional Pex3 and YFP-PTS1 co-localized punctae in these cells suggesting some peroxisomal remnants capable of matrix protein internalization (**Figure 6C**). In the *ABCD1^R518Q^* line, there was also a striking reduction in peroxisomal punctae, although here the remnant punctae were larger, clustered and more pronounced (**Figure 6D**). We repeated this analysis on flies at 25^0^C to examine the effect with a lower expression level. We observed that the human *ABCD1^Ref^* still led to a dramatic total loss of peroxisomes (**Figure 6E**). However, in the *ABCD1^R518Q^* line at the lower temperature we observed salivary gland peroxisomal punctae which were Pex3 and YFP positive suggesting only a partial impact on peroxisome biogenesis (**Figure 6F**). Taken together, these data suggest that overexpression of the human ABCD1 protein acts as a partial inhibitor of some aspect of peroxisomal biogenesis, or alternatively impacts degradation. s

**Figure 6.**
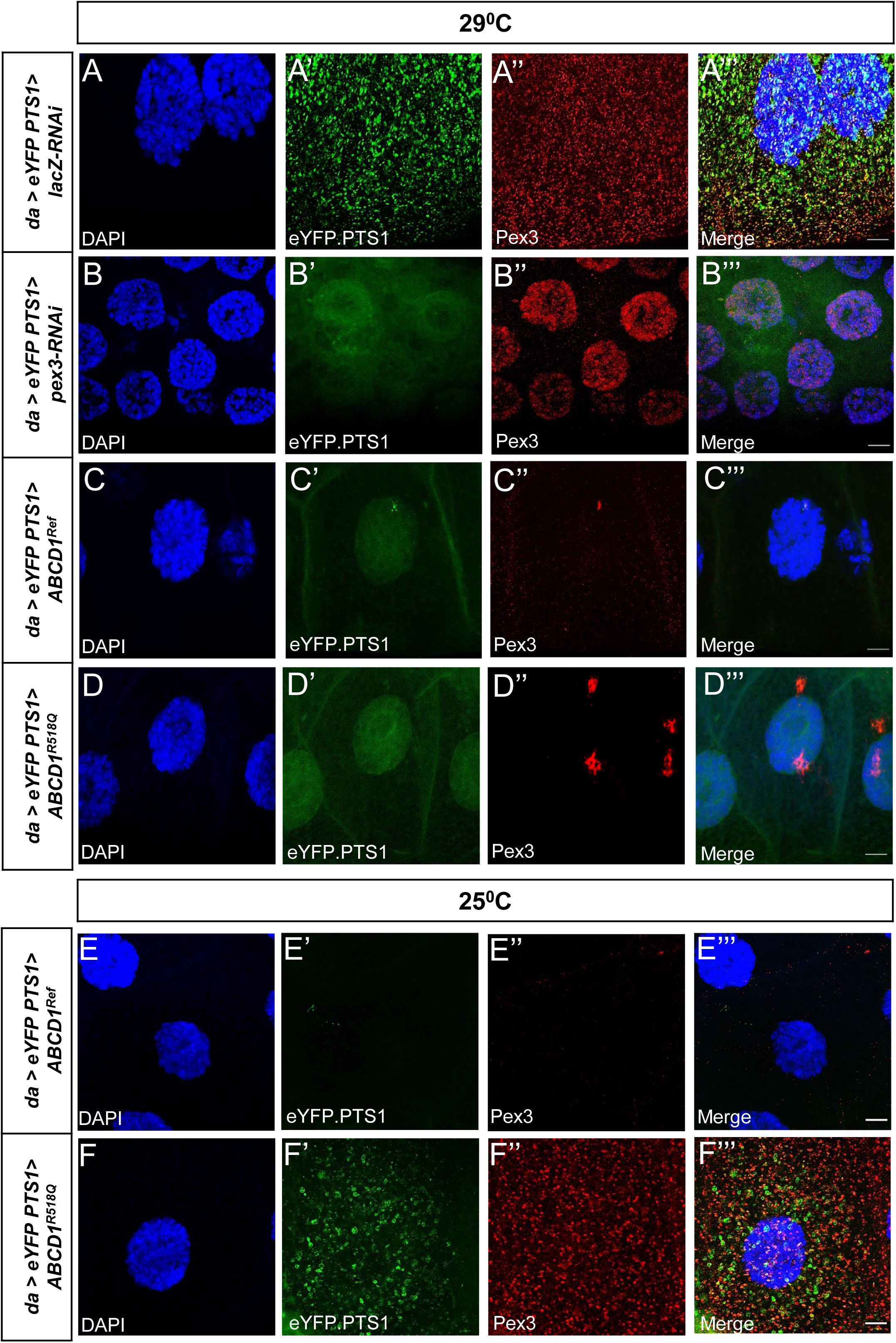
Peroxisomal biogenesis appeared to be affected in ubiquitous overexpression of human *ABCD1^Ref^* and *ABCD1^R518Q^* in larval salivary glands. *da-GAL4* is used to overexpress the peroxisomal marker UAS-*eYFP.PTS1* and the appropriate UAS lines at 29^0^C (**A-D**) and at 25^0^C (**E, F**). Anti-Pex3 antibody is also used to identify the peroxisomes. (**A**) *lacZ-RNAi* overexpression showed normal peroxisomal biogenesis and distribution per salivary gland cell. (**B**) *pex3-RNAi* overexpression resulted in the loss of peroxisomes along with a reduction in the salivary gland cell size. (**C**) Overexpression of *ABCD1^Ref^*and (**D**) *ABCD1^R528Q^* at 29^0^C led to a significant reduction in the number of peroxisomes. At 25^0^C, Overexpression of *ABCD1^Ref^* (**E**) similarly affects the peroxisomes, but the variant *ABCD1^R528Q^* (**F**) shows Pex3 positive peroxisomal punctae which are colocalizing with the YFP.PTS1 peroxisomal label. The Scale bar = 10µm.

To further confirm that functioning peroxisomes are reduced in the overexpression of ABCD1, we used LCMSMS in these larvae. This biochemical analysis showed an increase of 7.53-fold of C26:0 in *ABCD1^Ref^* as compared to *yw* control (**Figure 7A**), and 19.2-fold increase compared to *da>LacZ* control. Overexpressing the LOF variant *ABCD1^R518Q^* caused a milder and non-statistically significant accumulation of C26:0 by only 1.01-to-2.6-fold compared to control (**Figure 7A**). Using GCMS, which is calculated per larvae rather than lipid amount (**Figure 7B**), we also observed significant increases in C26:0 in both *ABCD1^Ref^* overexpression and *ABCD1^R518Q^* although in this case the mutant allele had a nominally higher value. Furthermore, *ABCD1^Ref^* showed significant increases of 140% in the ratios of C26:0/C22:0 over the control strains (**Figure 7C**), with *ABCD1^R518Q^*similarly showing a nominally higher value. No accumulation of C26:0-LPC was seen in *ABCD1^Ref^* larvae (**Supplementary figure 5**). When combining C26:0, C24:0 and C22:0 together, *ABCD1^Ref^* larvae showed about 3-fold accumulation compared to no accumulation in *ABCD1^R518Q^*but this trend did not reach statistical significance. Taken together the overexpression of human ABCD1 leads to both biochemical and morphological peroxisomal loss.

**Figure 7.**
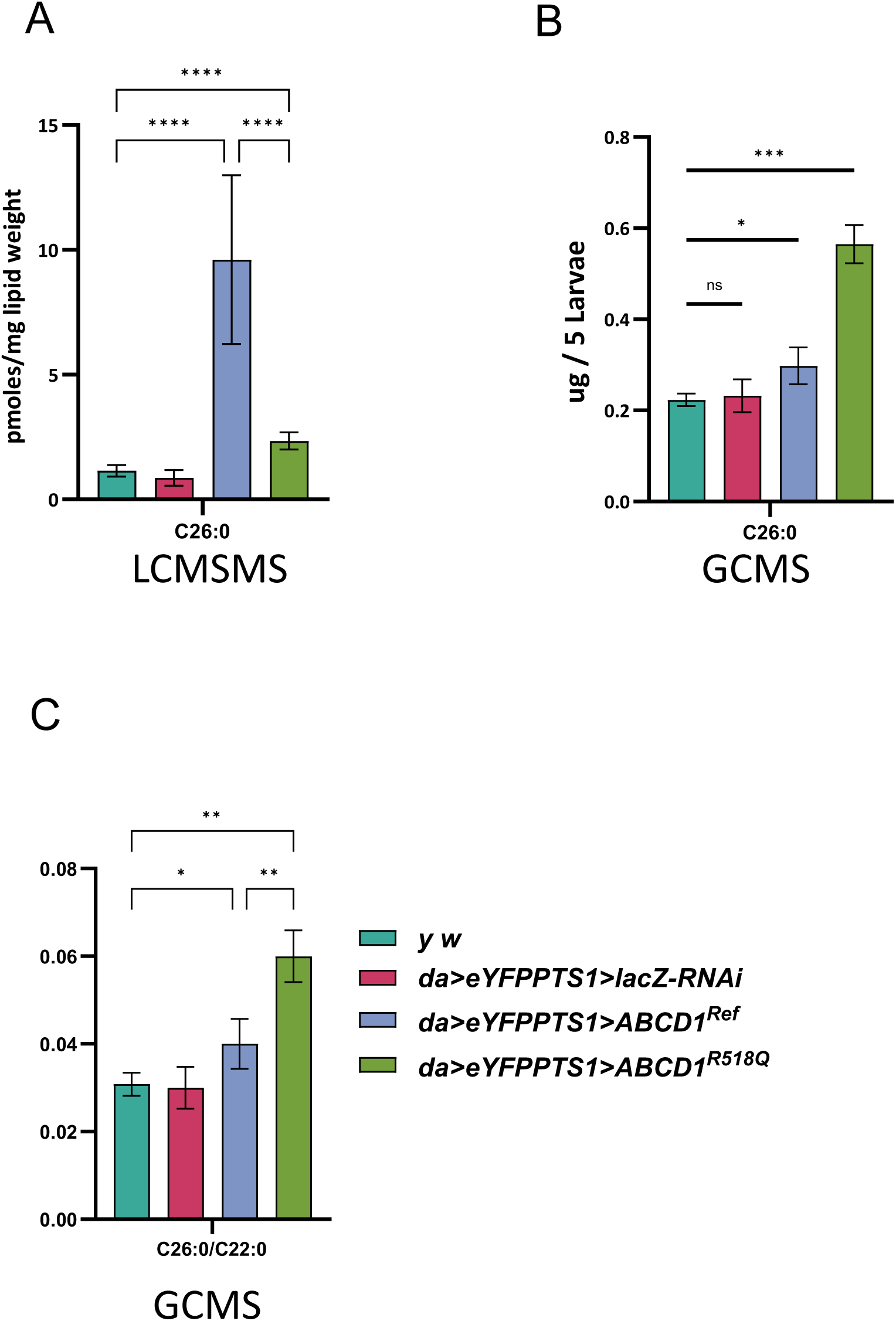
VLCFA analysis displayed elevated accumulation in human *ABCD1^Ref^* overexpression flies. A significant increase in the accumulation of C26:0 was observed in flies expressing the human *ABCD1^Ref^* in contrast to very mild accumulation of C26:0 in *ABCD1^R518Q^* in LCMS (**A**) and GCMS (**B**), see text for discussion about the differences. (**C**) Ratio of C26:0 to C22:0 VLCFA (as measured by GCMS) was observed to be elevated in *ABCD1^Ref^ and ABCD1^R518Q^*larvae compared to *yw* or da>lacZ control animals.

## DISCUSSION

*Drosophila* studies of X-ALD hold promise for providing novel insights into the molecular and biochemical pathogenesis of this neurological disorder. Some insight into X-ALD has come from studies on two VLCFA synthetases, *bgm* and *dbb.* LOF of *bgm* causes retinal degeneration, with the formation of vacuoles and the loss of photoreceptors and surrounding pigment cells. *dbb* knockout showed reduced survival and decreased locomotion activity and the accumulation of C26:1/C19:0, and C24:1/C19:0, fatty acids.^30^ For *bgm* LOF, both accumulation of VLCFA and degenerative symptoms were ameliorated with food enriched with oleic acid (C18:1),^42^ or with medium chain fatty acids.^29^ However these fly models are within the same VLCFA pathway but do not address the ortholog of the human *ABCD1* gene. Furthermore, the human orthologs of *bgm*, *ACSBG1* and *ACSBG2*, are not currently associated with any human disease.^43^ We also studied the *Acox1* gene in *Drosophila* downstream of *ABCD1,* and identified a unique *ACOX1* variant that led to gain-of-function and lipid droplet accumulation (Mitchell syndrome, MIM 618960).^28^ However, again this study does not address the *Drosophila Abcd1* ortholog. Although X-ALD is the most common peroxisomal disorder, *Drosophila* studies can provide mechanistic disease insights. Previous studies of *Drosophila Abcd1* have been somewhat limited by the location of the gene in the fly genome on the small heterochromatic 4^th^ chromosome, a chromosome in flies that has fewer genetic tools.^44^ This has limited the genetic studies of this gene in flies. A retinal phenotype was reported with RNAi knockdown of *Abcd1,* the only known study on this gene to date.^29^

Here, using RNAi knockdown and genetic mutation of *Abcd1*, we show that LOF of *Abcd1* impacts survival, motor function, the accumulation of the semeiotic C26:0, and buildup of lipid droplets (LD) in the retina. To the best of our knowledge, this is the first evidence of C26:0 accumulation in fly model of X-ALD. Our RNAi model, in contrast to the previous model^29^ that exhibited retinal degeneration, also demonstrates significantly reduced survival and peroxisomal abundance. The previous study used a different RNAi line *Abcd*^*HMS02382*^ and we utilized the *daughterless (da)* driver, a transcription factor expressed early in development, instead of the tubulin- and *repo-GAL4* drivers used in the earlier model.^29^ This difference in genetic drivers might explain the more pronounced developmental phenotype observed in our study. In our model, *Abcd1* was knocked down in photoreceptors using the *Rh1-GAL4* driver, and we observed the accumulation of interommatidial LD. We suspect that this LD accumulation is linked to the overproduction of reactive oxygen species (ROS). In humans and mice, *ABCD1* is primarily expressed in glial cells and is largely absent from neurons. In flies, LD accumulation in glia has been closely associated with ROS overproduction.^45,46^ The build-up of glial LDs has been connected to deficiency in *Acox1*.^28^ *ACOX1* deficiency is linked to a severe disorder that mimics peroxisomal biogenesis defects, known as pseudoneonatal adrenoleukodystrophy. Our results suggest a primarily glial metabolic alteration that could contribute to neurodegenerative pathology independently of neurons, similar to what is observed in *ACOX1* disorders.

In our model, elevations in VLCFAs were observed in both RNAi-mediated knockdown and genetic LOF of *Abcd1*. Despite lacking myelin, VLCFA are found in the form of ceramide phosphatidylethanolamine,^43^ which is thought to play a role in axonal wrapping and can lead to axonal dysfunction when elevated due to *Acox1* deficiency.^28^ Accumulation of VLCFAs may lead to increased production of sphingosine-1-phosphate (S1P) in glial cells,^47^ with excessive levels of S1P associated with a proinflammatory state and heightened production of ROS. S1P also has potent anti-apoptotic properties,^48^ and its overproduction could interfere with programmed cell death processes induced by ecdysone, potentially leading to the prolonged larval stage observed in our RNAi model. Thus, we suggest that *Abcd1* is involved in neurodegenerative mechanisms related to VLCFA toxicity, similarly to the synthetases and oxidases of VLCFA metabolism. We also observed a differential biochemical impact with a modest 50% decrease in *Abcd1* RNA which led to a 56-87% increase in C26:0-carnitine levels (and 51-72% in C26:0/C22:0 ratio) compared to both controls (*yw* and da>Luc). There was no impact on survival, fertility or motor function, or salivary gland morphology.

When the RNA reduction reached 90% in *da>Abcd1-RNAi^GD^*, we observed a severe phenotype characterized by significantly reduced survival, a 194-232% rise in C26:0 and 347-374% in C26:1(n-9). These flies also exhibited accumulation of C22:0, C24:0 and C24:1(n-9), in addition to abnormal ratio of C26:0/C22:0. A noticeable atrophy of the salivary gland and an impact on the number of peroxisomes per area was present. Additionally, LD accumulation in the retina was observed when expressed in photoreceptors, alongside evidence of abnormal eye formation with the eyeless driver seen in is a subset of flies – an unusual rosy eye appearance, echoing Pex16 deficiency (**Supplementary Figure 8**).^49^.

We also observed locomotor abnormalities in the *Abcd1*^Δ*4*^ flies, more noticeably in females. The *Abcd1*^Δ*4*^ demonstrated a dose-responsive accumulation of C26:0 – 373% compared to the control for *Abcd1*^Δ*4/*Δ*4*^ and 227% for *Abcd1*^Δ*4*/+^ (higher than the RNAi).

We have noted differences in survival and fertility rates between the RNAi knockdown and our Δ4 variant. Our initial results also show biochemical similarity in the levels of VLCFA in the tested larvae for RNAi and Δ4. Additional alleles of *Abcd1* may be needed to evaluate the Δ4 allele and whether it is truly a null allele. In addition, RNAi line can have off target effects, but our flies have VLCFA defects indicating that some of the phenotypes are likely due to the peroxisomal function.

In humans, levels of both C26:0-carnitine and C26:0-LPC are significantly elevated in ALD, yet in our fly model, we observed no such increase in LPC species. This discrepancy may be due to differences in phospholipase specificity that affects C26:0 hydrolysis. Invertebrates, including flies, lack homologs of some of the cytosolic phospholipase A2 (cPLA2)^50^ which in humans have a higher affinity for longer-chain fatty acids, such as C26:0^51^. Additionally, the availability of VLCFAs for hydrolysis is reduced by to their incorporation into LD, and in peroxisomal defects, LD can become resistant to lipolysis.^51^ Abundant LD formation are observed in our current model and previous models^28^ may further limit the conversion of VLCFA into LPC species, and further expand the interspecies differences in the observed VLCFA-LPC profile. Although overexpression of *ABCD1^Ref^* leads to the displacement of peroxisomes, only C26:0-carnitine and the combined group of (C22:0 + C24:0 + C26:0) acylcarnitine showed elevated levels, with other VLCFAs not showing similar increases. Lastly, our model showed elevations in C26:0 in both the RNAi and *Abcd1*^Δ*4*^ models; however, GCMS measurements did not indicate increases in the saturated VLCFA (C22, C24, and C26). Although GCMS is capable of detecting a wide range of lipids, it lacks the sensitivity of LCMS^52^. Specifically, for the non-volatile VLCFA, factors such as low ionization efficiency, peak broadening, and substrate thermolability contribute to reduced GCMS sensitivity compared to LCMS.^53–56^ In our methodology, only three parameters were measured using LCMS: C26:0, the sum of C22:0 + C24:0 + C26:0, and total VLCFA, limiting the extent of further comparisons.

The most surprising result of our work towards understanding ALD in flies is that we uncovered a dramatic effect of over-expression of human *ABCD1* in flies, namely an impact on peroxisome biogenesis itself. This was suspected based on a crumbled wing phenotype, resembling *pex2* mutants.^27^ Indeed, we saw nuclear staining of YFP-PTS1, similar to Pex3 RNAi knockdown,^26^ a phenotype not as strong as when overexpressing X-ALD associated *ABCD1^R518Q^*.

Genetic disorders of the peroxisome are typically divided into biogenesis disorders that disrupt the assembly of peroxisomes and single enzyme/protein defects, of which X-ALD is by far the most common. Conceptually, the peroxisome biogenesis machinery assembles the membrane and localizes the proteins while peroxisome proteins perform transport and catalyze the biochemistry necessary for peroxisome function. Peroxisomal single enzymes and proteins rely on PEX proteins for proper localization to peroxisomes. For example, PEX19 acts as a key chaperone for peroxisomal membrane proteins, such as ABCD1. ^57^ It is likely that overexpression of human ABCD1 in Drosophila strains overwhelms the capacity of endogenous peroxisome biogenesis proteins, like Pex19, to localize peroxisomal membrane proteins effectively. Another possibility is increased degradation or pexophagy in the cells expressing ABCD1. However, this seems less probable, as the ABCD1^R518Q^ construct at lower temperatures exhibits an intermediate phenotype characterized by larger, but not fewer, peroxisomes. Interestingly, although the phenotype in salivary glands resembled that of *Pex3* RNAi knockdown, we occasionally observed peroxisomal puncta capable of internalizing the YFP-PTS1 marker, indicating that some peroxisome assembly still occurs. Moreover, the ABCD1^R518Q^ variant line exhibited more of these peroxisomal puncta with intense anti-Pex3 staining. By lowering the temperature and reducing transgene expression, we observed an increase in the number of visible puncta. This data suggests that the peroxisome biogenesis machinery remains functional but is inhibited or overwhelmed by the presence of wild-type ABCD1. Most interestingly, a LOF variant appears to have a reduced inhibitory effect, especially at lower temperatures. These constructs, therefore, represent valuable new tools for studying peroxisome formation in Drosophila models. It will be interesting to express these human ABCD1 constructs in different temporal and spatial contexts in flies to fully assess the role of peroxisomes in a variety of biological contexts.

## CONCLUSION

Our results expand on the current model for peroxisomal defects associated with VLCFA esterification, their import into peroxisomes, and oxidation axis in the fruit fly. LOF of *Abcd1* results in a similar phenotype of reduced survival, retinal neurodegeneration, and locomotive impairment, and the accumulation of the pathognomonic C26:0. We show here for the first time the implication of excessive expression of the transporter, raising a suspicion for a dose-dependent pathology. Further research is needed to unveil the role of *ABCD1* LOF in early development of CNS pathology, and our results support the use of *Drosophila* to this end.

## ACKNOWLEDGEMENTS

We express our gratitude to Drs. Hugo Bellen, Shinya Yamamoto, Paul Marcogliese, and Matthew Moulton for their scientific advice and technical support. Our thanks also extend to Dr. Juan Botas for sharing his negative geotaxis robotic system for our research, and to Dr. Joshua Shulman for making his ERG equipment available to us. We are grateful to Drs. Hugo Bellen and James for generously donating fly stocks. Most importantly, we are deeply thankful to our patients and their families, who serve as a daily inspiration in our quest to discover new methods to enhance our care.

## FUNDING

This work was supported by the National Institute for Neurological Disorders and Stroke 5R01NS107733 to MFW.

### List of abbreviations

ABCD1 (hABCD1): human ATP binding cassette subfamily D member 1
Abcd1 (dABCD): *Drosophila melanogaster* ATP binding cassette subfamily D member 1
ABCD2: human ATP binding cassette subfamily D member 2
ALD: adrenoleukodystrophy
AMN: adrenomyeloneuropathy
ccALD: childhood cerebral ALD
CNS: central nervous system
CoA-VLCFA: very long chain fatty acids esterified to Co-enzyme A.
Da: daughterless
ERG: electroretinogram
GCMS: gas chromatography in tandem with mass spectrometry
GOF: gain of function
LCMS: high pressure liquid chromatography in tandem mass spectrometry
LD: lipid droplets
LOF: loss of function
LPC: Lysophosphatidylcholine
Luc: luciferase
NBD: nucleotide binding domain
NMD: nonsense mediated decay
PBS: phosphate-buffered saline
PCR: polymerase chain reaction
Rh1: neither inactivation nor afterpotential E - Rhodopsin 1
RNAi: Ribonucleic acid interference
SD: standard deviation
SEM: standard error of the mean
SKL: peroxisomal targeting sequence
TM: transmembrane domain
VLCFA: very long chain fatty acids
VUS: variant of uncertain significance
X-ALD: X-linked adrenoleukodystrophy

## REFERENCES

1. Engelen M, Kemp S, de Visser M, van Geel BM, Wanders RJ, Aubourg P, Poll-The BT. X-linked adrenoleukodystrophy (X-ALD): clinical presentation and guidelines for diagnosis, follow-up and management. Orphanet journal of rare diseases. 2012;7:51. Epub 2012/08/15. doi: 10.1186/1750-1172-7-51. PubMed PMID: 22889154; PMCID: Pmc3503704.

2. Moser AB, Jones RO, Hubbard WC, Tortorelli S, Orsini JJ, Caggana M, Vogel BH, Raymond GV. Newborn Screening for X-Linked Adrenoleukodystrophy. International journal of neonatal screening. 2016;2(4). Epub 2016/12/01. doi: 10.3390/ijns2040015. PubMed PMID: 31467997; PMCID: Pmc6715319.

3. Turk BR, Theda C, Fatemi A, Moser AB. X-linked adrenoleukodystrophy: Pathology, pathophysiology, diagnostic testing, newborn screening and therapies. International journal of developmental neuroscience : the official journal of the International Society for Developmental Neuroscience. 2020;80(1):52–72. Epub 2020/01/08. doi: 10.1002/jdn.10003. PubMed PMID: 31909500; PMCID: Pmc7041623.

4. Weinhofer I, Zierfuss B, Hametner S, Wagner M, Popitsch N, Machacek C, Bartolini B, Zlabinger G, Ohradanova-Repic A, Stockinger H, Köhler W, Höftberger R, Regelsberger G, Forss-Petter S, Lassmann H, Berger J. Impaired plasticity of macrophages in X-linked adrenoleukodystrophy. Brain : a journal of neurology. 2018;141(8):2329–42. doi: 10.1093/brain/awy127. PubMed PMID: 29860501; PMCID: PMC6061697.

5. Manor J, Chung H, Bhagwat PK, Wangler MF. ABCD1 and X-linked adrenoleukodystrophy: A disease with a markedly variable phenotype showing conserved neurobiology in animal models. Journal of neuroscience research. 2021;99(12):3170–81. Epub 2021/10/31. doi: 10.1002/jnr.24953. PubMed PMID: 34716609.

6. Xiong C, Jia LN, Xiong WX, Wu XT, Xiong LL, Wang TH, Zhou D, Hong Z, Liu Z, Tang L. Structural insights into substrate recognition and translocation of human peroxisomal ABC transporter ALDP. Signal transduction and targeted therapy. 2023;8(1):74. Epub 20230222. doi: 10.1038/s41392-022-01280-9. PubMed PMID: 36810450; PMCID: PMC9944889.

7. Mallack EJ, Gao K, Engelen M, Kemp S. Structure and Function of the ABCD1 Variant Database: 20 Years, 940 Pathogenic Variants, and 3400 Cases of Adrenoleukodystrophy. Cells. 2022;11(2). Epub 20220114. doi: 10.3390/cells11020283. PubMed PMID: 35053399; PMCID: PMC8773697.

8. Wiesinger C, Eichler FS, Berger J. The genetic landscape of X-linked adrenoleukodystrophy: inheritance, mutations, modifier genes, and diagnosis. Appl Clin Genet. 2015;8:109–21. Epub 20150502. doi: 10.2147/tacg.S49590. PubMed PMID: 25999754; PMCID: PMC4427263.

9. Kemp S, Pujol A, Waterham HR, van Geel BM, Boehm CD, Raymond GV, Cutting GR, Wanders RJ, Moser HW. ABCD1 mutations and the X-linked adrenoleukodystrophy mutation database: role in diagnosis and clinical correlations. Human mutation. 2001;18(6):499–515. doi: 10.1002/humu.1227. PubMed PMID: 11748843.

10. Berger J, Forss-Petter S, Eichler FS. Pathophysiology of X-linked adrenoleukodystrophy. Biochimie. 2014;98(100):135–42. Epub 2013/12/10. doi: 10.1016/j.biochi.2013.11.023. PubMed PMID: 24316281; PMCID: Pmc3988840.

11. Höftberger R, Kunze M, Weinhofer I, Aboul-Enein F, Voigtländer T, Oezen I, Amann G, Bernheimer H, Budka H, Berger J. Distribution and cellular localization of adrenoleukodystrophy protein in human tissues: implications for X-linked adrenoleukodystrophy. Neurobiology of disease. 2007;28(2):165–74. Epub 2007/09/01. doi: 10.1016/j.nbd.2007.07.007. PubMed PMID: 17761426.

12. Schackmann MJ, Ofman R, Dijkstra IM, Wanders RJ, Kemp S. Enzymatic characterization of ELOVL1, a key enzyme in very long-chain fatty acid synthesis. Biochimica et biophysica acta. 2015;1851(2):231–7. Epub 20141211. doi: 10.1016/j.bbalip.2014.12.005. PubMed PMID: 25499606.

13. Zierfuss B, Buda A, Villoria-González A, Logist M, Fabjan J, Parzer P, Battin C, Vandersteene S, Dijkstra IME, Waidhofer-Söllner P, Grabmeier-Pfistershammer K, Steinberger P, Kemp S, Forss-Petter S, Berger J, Weinhofer I. Saturated very long-chain fatty acids regulate macrophage plasticity and invasiveness. J Neuroinflammation. 2022;19(1):305. Epub 20221217. doi: 10.1186/s12974-022-02664-y. PubMed PMID: 36528616; PMCID: PMC9759912.

14. Buda A, Forss-Petter S, Hua R, Jaspers Y, Lassnig M, Waidhofer-Söllner P, Kemp S, Kim P, Weinhofer I, Berger J. ABCD1 Transporter Deficiency Results in Altered Cholesterol Homeostasis. Biomolecules. 2023;13(9). Epub 20230831. doi: 10.3390/biom13091333. PubMed PMID: 37759733; PMCID: PMC10526550.

15. Singh I, Pujol A. Pathomechanisms underlying X-adrenoleukodystrophy: a three-hit hypothesis. Brain pathology (Zurich, Switzerland). 2010;20(4):838–44. doi: 10.1111/j.1750-3639.2010.00392.x. PubMed PMID: 20626745; PMCID: PMC3021280.

16. Pujol A, Ferrer I, Camps C, Metzger E, Hindelang C, Callizot N, Ruiz M, Pàmpols T, Giròs M, Mandel JL. Functional overlap between ABCD1 (ALD) and ABCD2 (ALDR) transporters: a therapeutic target for X-adrenoleukodystrophy. Human molecular genetics. 2004;13(23):2997–3006. Epub 2004/10/19. doi: 10.1093/hmg/ddh323. PubMed PMID: 15489218.

17. Muneer Z, Wiesinger C, Voigtländer T, Werner HB, Berger J, Forss-Petter S. Abcd2 is a strong modifier of the metabolic impairments in peritoneal macrophages of ABCD1-deficient mice. PloS one. 2014;9(9):e108655. Epub 2014/09/26. doi: 10.1371/journal.pone.0108655. PubMed PMID: 25255441; PMCID: Pmc4177892.

18. Matsukawa T, Asheuer M, Takahashi Y, Goto J, Suzuki Y, Shimozawa N, Takano H, Onodera O, Nishizawa M, Aubourg P, Tsuji S. Identification of novel SNPs of ABCD1, ABCD2, ABCD3, and ABCD4 genes in patients with X-linked adrenoleukodystrophy (ALD) based on comprehensive resequencing and association studies with ALD phenotypes. Neurogenetics. 2011;12(1):41–50. Epub 2010/07/28. doi: 10.1007/s10048-010-0253-6. PubMed PMID: 20661612; PMCID: Pmc3029816.

19. Martinović K, Bauer J, Kunze M, Berger J, Forss-Petter S. Abcd1 deficiency accelerates cuprizone-induced oligodendrocyte loss and axonopathy in a demyelinating mouse model of X-linked adrenoleukodystrophy. Acta Neuropathol Commun. 2023;11(1):98. Epub 20230618. doi: 10.1186/s40478-023-01595-w. PubMed PMID: 37331971; PMCID: PMC10276915.

20. Pujol A, Hindelang C, Callizot N, Bartsch U, Schachner M, Mandel JL. Late onset neurological phenotype of the X-ALD gene inactivation in mice: a mouse model for adrenomyeloneuropathy. Human molecular genetics. 2002;11(5):499–505. doi: 10.1093/hmg/11.5.499. PubMed PMID: 11875044.

21. McGuinness MC, Lu JF, Zhang HP, Dong GX, Heinzer AK, Watkins PA, Powers J, Smith KD. Role of ALDP (ABCD1) and mitochondria in X-linked adrenoleukodystrophy. Molecular and cellular biology. 2003;23(2):744–53. doi: 10.1128/mcb.23.2.744-753.2003. PubMed PMID: 12509471; PMCID: PMC151532.

22. Wiesinger C, Kunze M, Regelsberger G, Forss-Petter S, Berger J. Impaired very long-chain acyl-CoA β-oxidation in human X-linked adrenoleukodystrophy fibroblasts is a direct consequence of ABCD1 transporter dysfunction. The Journal of biological chemistry. 2013;288(26):19269–79. Epub 20130513. doi: 10.1074/jbc.M112.445445. PubMed PMID: 23671276; PMCID: PMC3696697.

23. Yamamoto S, Kanca O, Wangler MF, Bellen HJ. Integrating non-mammalian model organisms in the diagnosis of rare genetic diseases in humans. Nature reviews Genetics. 2024;25(1):46–60. Epub 20230725. doi: 10.1038/s41576-023-00633-6. PubMed PMID: 37491400.

24. Baldridge D, Wangler MF, Bowman AN, Yamamoto S, Schedl T, Pak SC, Postlethwait JH, Shin J, Solnica-Krezel L, Bellen HJ, Westerfield M. Model organisms contribute to diagnosis and discovery in the undiagnosed diseases network: current state and a future vision. Orphanet journal of rare diseases. 2021;16(1):206. Epub 20210507. doi: 10.1186/s13023-021-01839-9. PubMed PMID: 33962631; PMCID: PMC8103593.

25. Wangler MF, Chao YH, Roth M, Welti R, McNew JA. Drosophila Models Uncover Substrate Channeling Effects on Phospholipids and Sphingolipids in Peroxisomal Biogenesis Disorders. bioRxiv. 2024. Epub 20240429. doi: 10.1101/2024.04.26.591192. PubMed PMID: 38746221; PMCID: PMC11092477.

26. Wangler MF, Chao YH, Bayat V, Giagtzoglou N, Shinde AB, Putluri N, Coarfa C, Donti T, Graham BH, Faust JE, McNew JA, Moser A, Sardiello M, Baes M, Bellen HJ. Peroxisomal biogenesis is genetically and biochemically linked to carbohydrate metabolism in Drosophila and mouse. PLoS genetics. 2017;13(6):e1006825. Epub 20170622. doi: 10.1371/journal.pgen.1006825. PubMed PMID: 28640802; PMCID: PMC5480855.

27. Faust JE, Manisundaram A, Ivanova PT, Milne SB, Summerville JB, Brown HA, Wangler M, Stern M, McNew JA. Peroxisomes are required for lipid metabolism and muscle function in Drosophila melanogaster. PloS one. 2014;9(6):e100213. Epub 20140619. doi: 10.1371/journal.pone.0100213. PubMed PMID: 24945818; PMCID: PMC4063865.

28. Chung HL, Wangler MF, Marcogliese PC, Jo J, Ravenscroft TA, Zuo Z, Duraine L, Sadeghzadeh S, Li-Kroeger D, Schmidt RE, Pestronk A, Rosenfeld JA, Burrage L, Herndon MJ, Chen S, Shillington A, Vawter-Lee M, Hopkin R, Rodriguez-Smith J, Henrickson M, Lee B, Moser AB, Jones RO, Watkins P, Yoo T, Mar S, Choi M, Bucelli RC, Yamamoto S, Lee HK, Prada CE, Chae JH, Vogel TP, Bellen HJ. Loss- or Gain-of-Function Mutations in ACOX1 Cause Axonal Loss via Different Mechanisms. Neuron. 2020;106(4):589–606.e6. Epub 2020/03/15. doi: 10.1016/j.neuron.2020.02.021. PubMed PMID: 32169171; PMCID: Pmc7289150.

29. Gordon HB, Valdez L, Letsou A. Etiology and treatment of adrenoleukodystrophy: new insights from Drosophila. Disease models & mechanisms. 2018;11(6). Epub 2018/05/10. doi: 10.1242/dmm.031286. PubMed PMID: 29739804; PMCID: Pmc6031365.

30. Sivachenko A, Gordon HB, Kimball SS, Gavin EJ, Bonkowsky JL, Letsou A. Neurodegeneration in a Drosophila model of adrenoleukodystrophy: the roles of the Bubblegum and Double bubble acyl-CoA synthetases. Disease models & mechanisms. 2016;9(4):377–87. Epub 2016/02/20. doi: 10.1242/dmm.022244. PubMed PMID: 26893370; PMCID: Pmc4852500.

31. Untergasser A, Cutcutache I, Koressaar T, Ye J, Faircloth BC, Remm M, Rozen SG. Primer3--new capabilities and interfaces. Nucleic acids research. 2012;40(15):e115. Epub 20120622. doi: 10.1093/nar/gks596. PubMed PMID: 22730293; PMCID: PMC3424584.

32. Onur TS, Laitman A, Zhao H, Keyho R, Kim H, Wang J, Mair M, Wang H, Li L, Perez A, de Haro M, Wan YW, Allen G, Lu B, Al-Ramahi I, Liu Z, Botas J. Downregulation of glial genes involved in synaptic function mitigates Huntington’s disease pathogenesis. eLife. 2021;10. Epub 20210419. doi: 10.7554/eLife.64564. PubMed PMID: 33871358; PMCID: PMC8149125.

33. Verstreken P, Koh TW, Schulze KL, Zhai RG, Hiesinger PR, Zhou Y, Mehta SQ, Cao Y, Roos J, Bellen HJ. Synaptojanin is recruited by endophilin to promote synaptic vesicle uncoating. Neuron. 2003;40(4):733–48. doi: 10.1016/s0896-6273(03)00644-5. PubMed PMID: 14622578.

34. Chung HL, Augustine GJ, Choi KW. Drosophila Schip1 Links Expanded and Tao-1 to Regulate Hippo Signaling. Dev Cell. 2016;36(5):511–24. doi: 10.1016/j.devcel.2016.02.004. PubMed PMID: 26954546.

35. Falkenberg KD, Braverman NE, Moser AB, Steinberg SJ, Klouwer FCC, Schlüter A, Ruiz M, Pujol A, Engvall M, Naess K, van Spronsen F, Körver-Keularts I, Rubio-Gozalbo ME, Ferdinandusse S, Wanders RJA, Waterham HR. Allelic Expression Imbalance Promoting a Mutant PEX6 Allele Causes Zellweger Spectrum Disorder. American journal of human genetics. 2017;101(6):965–76. doi: 10.1016/j.ajhg.2017.11.007. PubMed PMID: 29220678; PMCID: PMC5812895.

36. Lagerstedt SA, Hinrichs DR, Batt SM, Magera MJ, Rinaldo P, McConnell JP. Quantitative determination of plasma c8–c26 total fatty acids for the biochemical diagnosis of nutritional and metabolic disorders. Mol Genet Metab. 2001;73(1):38–45.

37. van de Beek M-C, Dijkstra IM, van Lenthe H, Ofman R, Goldhaber-Pasillas D, Schauer N, Schackmann M, Engelen-Lee J-Y, Vaz FM, Kulik W. C26: 0-carnitine is a new biomarker for X-linked adrenoleukodystrophy in mice and man. PloS one. 2016;11(4):e0154597.

38. Sayers EW, Bolton EE, Brister JR, Canese K, Chan J, Comeau DC, Connor R, Funk K, Kelly C, Kim S, Madej T, Marchler-Bauer A, Lanczycki C, Lathrop S, Lu Z, Thibaud-Nissen F, Murphy T, Phan L, Skripchenko Y, Tse T, Wang J, Williams R, Trawick BW, Pruitt KD, Sherry ST. Database resources of the national center for biotechnology information. Nucleic acids research. 2022;50(D1):D20-d6. doi: 10.1093/nar/gkab1112. PubMed PMID: 34850941; PMCID: PMC8728269.

39. Caudy M, Vässin H, Brand M, Tuma R, Jan LY, Jan YN. daughterless, a Drosophila gene essential for both neurogenesis and sex determination, has sequence similarities to myc and the achaete-scute complex. Cell. 1988;55(6):1061–7. doi: 10.1016/0092-8674(88)90250-4. PubMed PMID: 3203380.

40. Engelen M, Kemp S, Poll-The BT. X-linked adrenoleukodystrophy: pathogenesis and treatment. Current neurology and neuroscience reports. 2014;14(10):486. Epub 2014/08/15. doi: 10.1007/s11910-014-0486-0. PubMed PMID: 25115486.

41. Rattay TW, Rautenberg M, Söhn AS, Hengel H, Traschütz A, Röben B, Hayer SN, Schüle R, Wiethoff S, Zeltner L, Haack TB, Cegan A, Schöls L, Schleicher E, Peter A. Defining diagnostic cutoffs in neurological patients for serum very long chain fatty acids (VLCFA) in genetically confirmed X-Adrenoleukodystrophy. Scientific reports. 2020;10(1):15093. Epub 20200915. doi: 10.1038/s41598-020-71248-8. PubMed PMID: 32934269; PMCID: PMC7494896.

42. Min KT, Benzer S. Preventing neurodegeneration in the Drosophila mutant bubblegum. Science (New York, NY). 1999;284(5422):1985-8. doi: 10.1126/science.284.5422.1985. PubMed PMID: 10373116.

43. Watkins PA. Fatty Acyl-CoA Synthetases. In: Lennarz WJ, Lane MD, editors. Encyclopedia of Biological Chemistry (Second Edition). Waltham: Academic Press; 2013. p. 290-5.

44. Öztürk-Çolak A, Marygold SJ, Antonazzo G, Attrill H, Goutte-Gattat D, Jenkins VK, Matthews BB, Millburn G, Dos Santos G, Tabone CJ. FlyBase: updates to the Drosophila genes and genomes database. Genetics. 2024;227(1). doi: 10.1093/genetics/iyad211. PubMed PMID: 38301657; PMCID: PMC11075543.

45. Liu L, Zhang K, Sandoval H, Yamamoto S, Jaiswal M, Sanz E, Li Z, Hui J, Graham BH, Quintana A, Bellen HJ. Glial lipid droplets and ROS induced by mitochondrial defects promote neurodegeneration. Cell. 2015;160(1-2):177–90. doi: 10.1016/j.cell.2014.12.019. PubMed PMID: 25594180; PMCID: PMC4377295.

46. Cabirol-Pol MJ, Khalil B, Rival T, Faivre-Sarrailh C, Besson MT. Glial lipid droplets and neurodegeneration in a Drosophila model of complex I deficiency. Glia. 2018;66(4):874–88. Epub 20171229. doi: 10.1002/glia.23290. PubMed PMID: 29285794.

47. Chung HL, Ye Q, Park YJ, Zuo Z, Mok JW, Kanca O, Tattikota SG, Lu S, Perrimon N, Lee HK, Bellen HJ. Very-long-chain fatty acids induce glial-derived sphingosine-1-phosphate synthesis, secretion, and neuroinflammation. Cell metabolism. 2023;35(5):855–74.e5. Epub 20230420. doi: 10.1016/j.cmet.2023.03.022. PubMed PMID: 37084732; PMCID: PMC10160010.

48. Phan VH, Herr DR, Panton D, Fyrst H, Saba JD, Harris GL. Disruption of sphingolipid metabolism elicits apoptosis-associated reproductive defects in Drosophila. Dev Biol. 2007;309(2):329–41. Epub 20070726. doi: 10.1016/j.ydbio.2007.07.021. PubMed PMID: 17706961; PMCID: PMC2094363.

49. Nakayama M, Sato H, Okuda T, Fujisawa N, Kono N, Arai H, Suzuki E, Umeda M, Ishikawa HO, Matsuno K. Drosophila carrying pex3 or pex16 mutations are models of Zellweger syndrome that reflect its symptoms associated with the absence of peroxisomes. PloS one. 2011;6(8):e22984. Epub 20110803. doi: 10.1371/journal.pone.0022984. PubMed PMID: 21826223; PMCID: PMC3149631.

50. Murakami M, Sato H, Taketomi Y. Updating Phospholipase A(2) Biology. Biomolecules. 2020;10(10). Epub 20201019. doi: 10.3390/biom10101457. PubMed PMID: 33086624; PMCID: PMC7603386.

51. Hayashi D, Mouchlis VD, Dennis EA. Omega-3 versus Omega-6 fatty acid availability is controlled by hydrophobic site geometries of phospholipase A(2)s. Journal of lipid research. 2021;62:100113. Epub 20210830. doi: 10.1016/j.jlr.2021.100113. PubMed PMID: 34474084; PMCID: PMC8551542.

52. Koch E, Wiebel M, Hopmann C, Kampschulte N, Schebb NH. Rapid quantification of fatty acids in plant oils and biological samples by LC-MS. Analytical and bioanalytical chemistry. 2021;413(21):5439–51. Epub 20210722. doi: 10.1007/s00216-021-03525-y. PubMed PMID: 34296318; PMCID: PMC8405509.

53. Wang D, Yu S, Zhang Y, Yin Y, Cheng Q, Xie S, Yu J, Li H, Cheng X, Qiu L. Rapid liquid chromatography-tandem mass spectrometry to determine very-long-chain fatty acids in human and to establish reference intervals for the Chinese population. Clinica chimica acta; international journal of clinical chemistry. 2019;495:185–90. Epub 20190409. doi: 10.1016/j.cca.2019.04.058. PubMed PMID: 30978326.

54. Wangler MF, Lesko B, Dahal R, Jangam S, Bhadane P, Wilson TE, McPheron M, Miller MJ. Dicarboxylic acylcarnitine biomarkers in peroxisome biogenesis disorders. Mol Genet Metab. 2023;140(3):107680. Epub 20230807. doi: 10.1016/j.ymgme.2023.107680. PubMed PMID: 37567036; PMCID: PMC10840807.

55. Seppänen-Laakso T, Laakso I, Hiltunen R. Analysis of fatty acids by gas chromatography, and its relevance to research on health and nutrition. Analytica Chimica Acta. 2002;465(1-2):39–62.

56. Hoving LR, Heijink M, van Harmelen V, van Dijk KW, Giera M. GC-MS analysis of medium-and long-chain fatty acids in blood samples. Clinical Metabolomics: Methods and Protocols. 2018:257–65.

57. Sacksteder KA, Jones JM, South ST, Li X, Liu Y, Gould SJ. PEX19 binds multiple peroxisomal membrane proteins, is predominantly cytoplasmic, and is required for peroxisome membrane synthesis. The Journal of cell biology. 2000;148(5):931–44. doi: 10.1083/jcb.148.5.931. PubMed PMID: 10704444; PMCID: PMC2174547.

