## Supplemental Materials for "Genetic analysis of the X-linked Adrenoleukodystrophy *ABCD1 gene* in *Drosophila* uncovers a role in Peroxisomal dynamics"

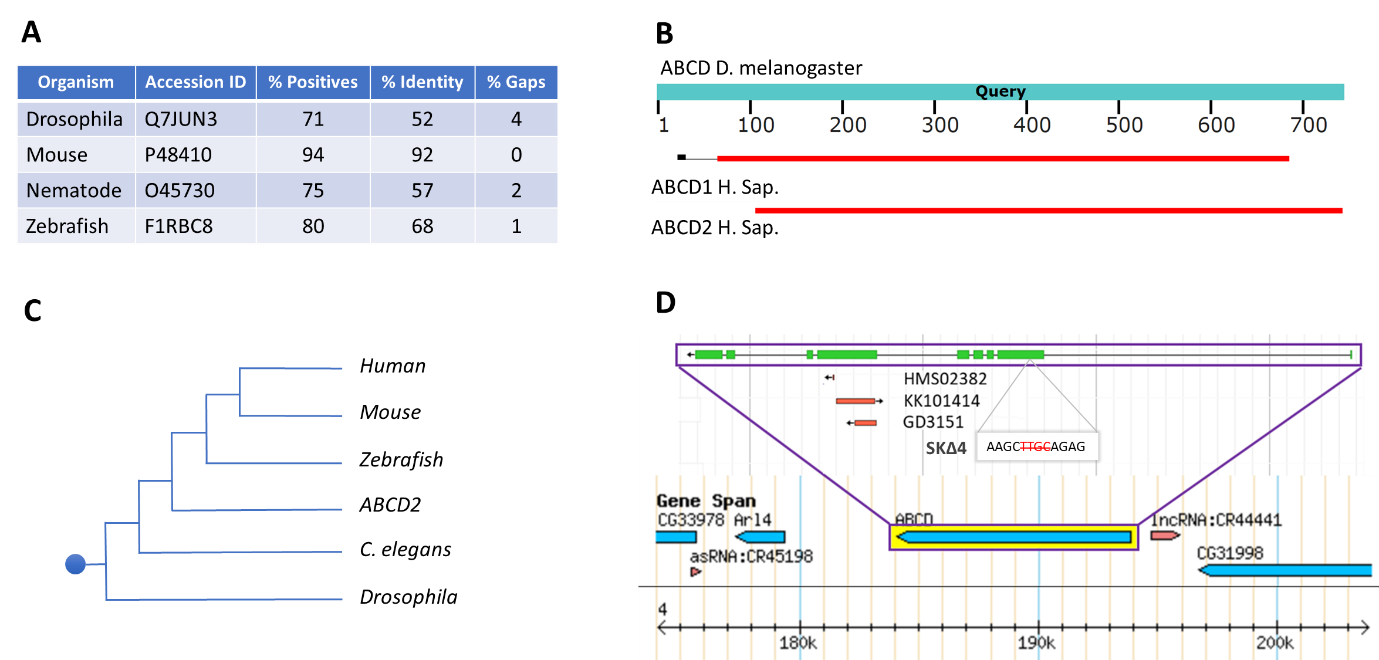

**Supplementary Figure 1.** Bioinformatics for ABCD1. (**A**) A table comparing homology of the human ABCD1 protein with orthologs for which animal models have been developed.^1-4^ Nematodes refer to %*Caenorhabditis elegans*. (**B**) An alignment of Drosophila’s *Abcd1* gene with human *ABCD1* and *ABCD2* genes, demonstrating high homology to both (and in between the human *ABCD1* and *ABCD2*). The alignment was conducted using BLAST.^5^ (**C**) A schematic illustration of a phylogenetic cladogram depicting the evolutionary relationships of *ABCD1* among various species (and with *ABCD2*). The scheme is based on Fourcade et. al.^6^ (**D**) A schematic depiction of the locus containing the Abcd1 gene on chromosome 4 (bottom panel) alongside the three stocks of dsRNA and the genomic out-of-frame deletion within the first exon utilized in this work (upper panel). The scheme was rendered using Flybase.org.

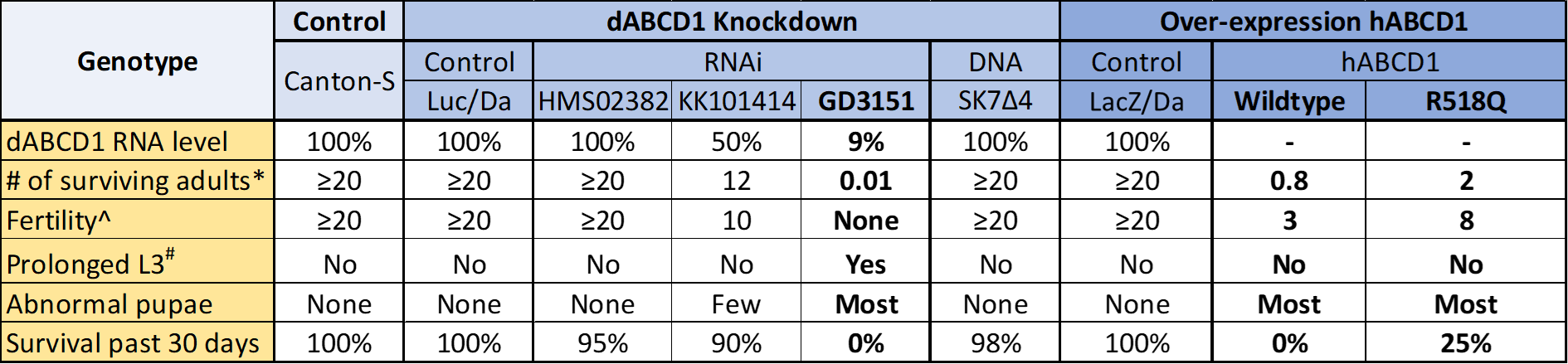

**Supplementary Table 1:** Comparison of Viability, Pupariation, and Fertility Among Control Flies and Flies with *Abcd1* Knockdown and Overexpression of Human *ABCD1*. * – Represents the average number of flies of the specified genotype emerging per 1 female parent (the offspring carries the specified genotype, while the parent may have a different genotype, depending on the cross). ^ – Denotes the number of offspring successfully emerging per female (the female parent has the specified genotype, though the offspring may have a different genotype, depending on the cross). # – Identifies larvae exhibiting a prolonged 3rd instar stage without undergoing pupariation. To increase clarity, Drosophila’s *Abcd1* is labeled here as dABCD1, whereas the human ABCD1 is referred to as hABCD1. R518Q – Indicates the c.1553G>A pathogenic variant inserted into the wildtype sequence of hABCD1. Controls include the Canton-S wildtype fly, UAS-Luciferase RNAi expressed under the da-Gal4 driver (Luc/Da), and UAS-LacZ overexpression under the da-Gal4 driver for hABCD1 overexpression (LacZ/Da).

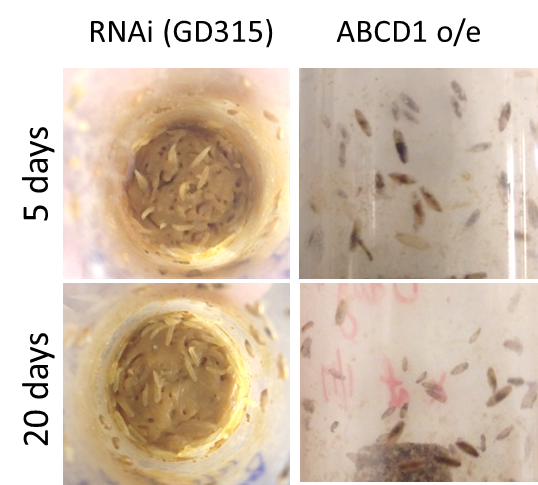

**Supplementary Figure 2:** Abnormal Offspring from Knockdown and Overexpression of Drosophila ABCD. The left panels display a representative image of a standard tube containing third instar larvae expressing RNAi against endogenous *Abcd1* (dsRNA strain GD3151) at 5 days (top) and 20 days (bottom) after their first appearance, showing a prolonged third instar stage without undergoing pupariation. On the right – Overexpression (o/e) of human *ABCD1* led to uniformly abnormal pupae (indicated by darker segments within the pupae).

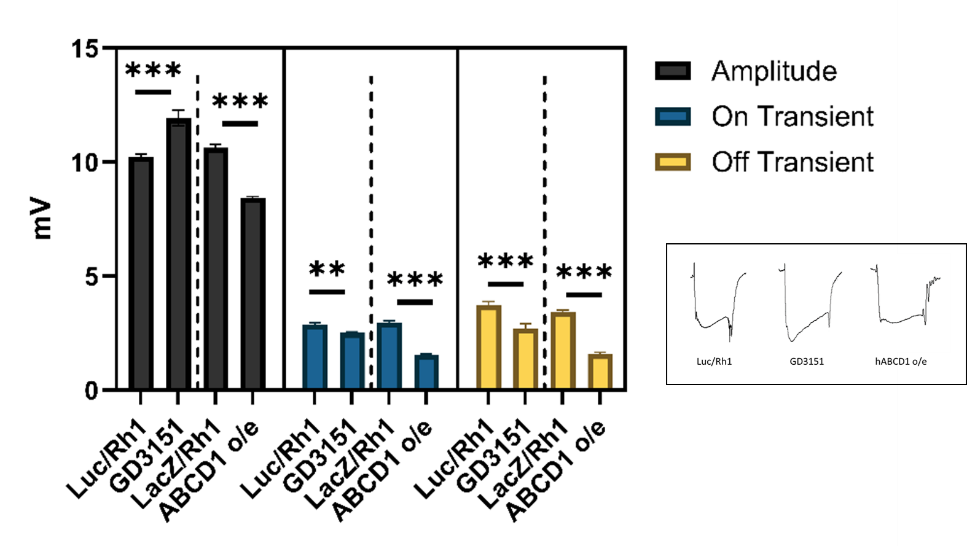

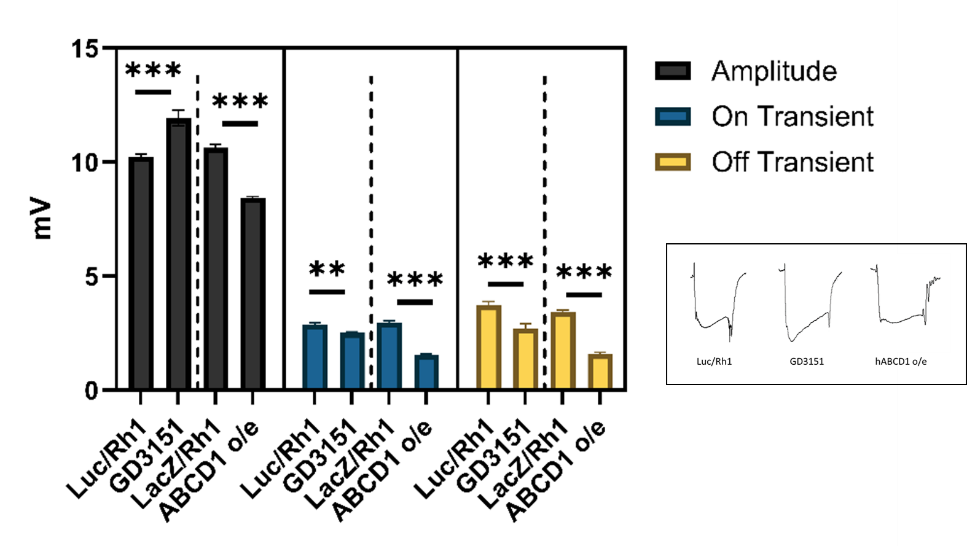

**Supplementary Figure 3**. **Electroretinogram assay in detail comparing *Abcd1* Knockdown and Human *ABCD1* overexpression** (to the right of the dashed line) with Controls. In the knockdown analysis, Luc/Rh1 serves as the control compared to *Abcd1* RNAi (Rh1/UAS-ABCD^GD3151^), using luciferase RNAi as a non-target control via the Rh1-Gal4 driver. For overexpression analysis, LacZ/Rh1, expressing LacZ using the same Rh1-Gal4 driver, acts as control. The average electroretinogram amplitude (gray bars) shows a 17% increase for the knockdown group (10.22 vs 11.925 mV), while there is a 21% decrease from control to human *ABCD1* overexpression (8.425 vs 10.625 mV). Moreover, the depolarizing "on transient" amplitude decreases by 13% in RNAi and by 48% in overexpression compared to their respective controls. Likewise, the depolarizing "off transient" amplitude reduces by 27% in RNAi and by 54% in overexpression cases. These differences reached statistical significance, with ** indicating a p-value <0.01 and *** indicating a p-value <0.001. No statistical difference was observed between Luc/Rh1 and LacZ/Rh1 across any of the three measured parameters. The data represents averages from 24 retinograms across 8 flies, with error bars showing the SEM (standard error of the mean). Statistical analysis was conducted using the Mann-Whitney U test for nonparametric data, due to the distribution not meeting all criteria for uniformity. Inset: representative retinograms for the control (Luc/Rh1), GD3151, and hABCD1 o/e cases.

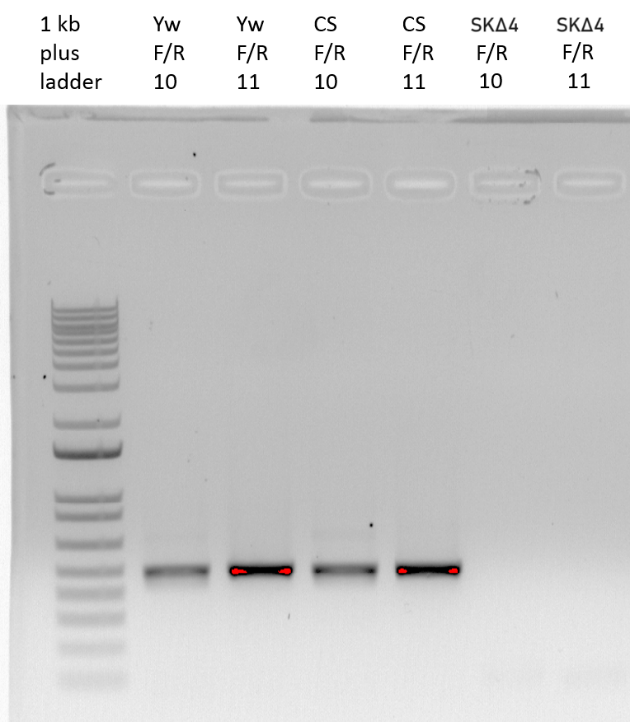

**Supplementary Figure 4**: **Confirmation of Genomic Out-of-Frame Deletion**. The 4nt deletion line (SKΔ4) failed to amplify the segment between the forward primer labeled as 10 and the reverse primer 11, the region where the deletion is located. The control flies *y1w1* *(yw)* and Canton-S (CS) successfully amplified this segment.

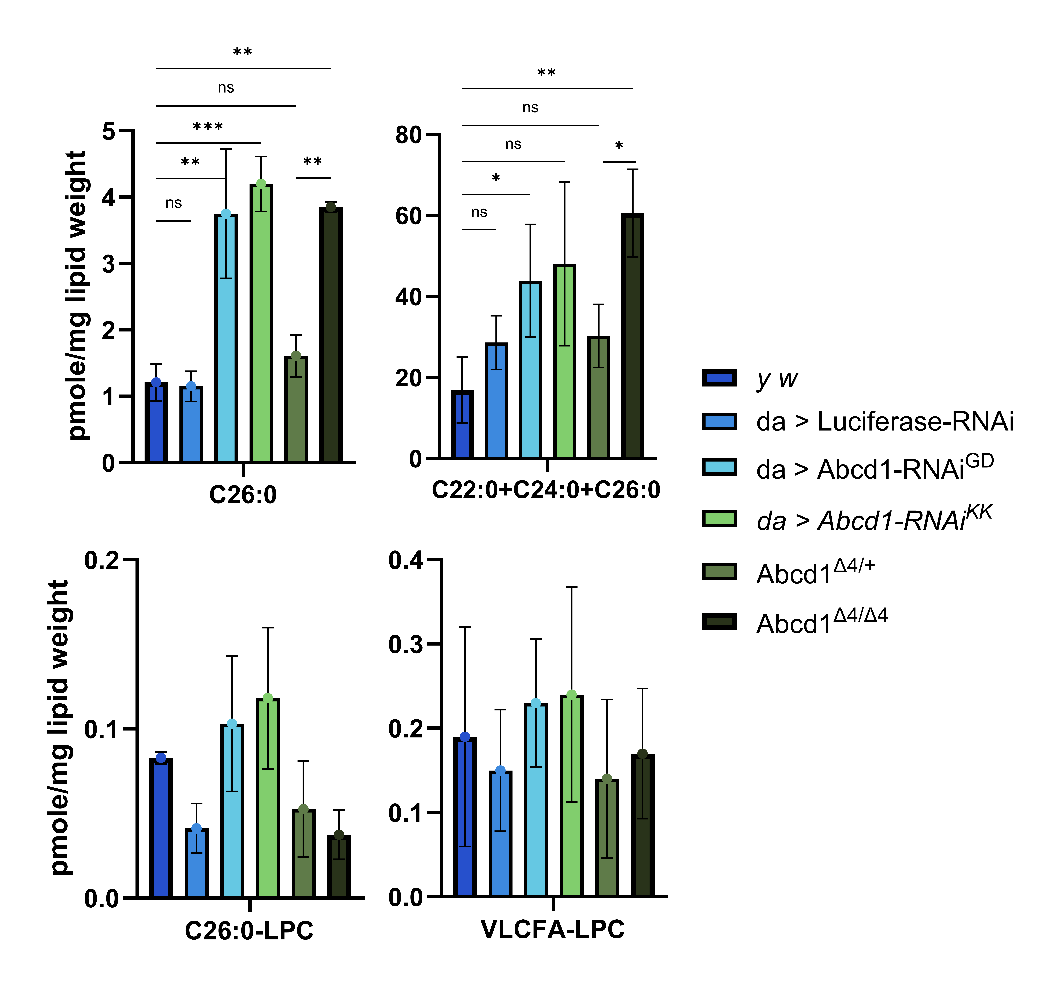

**Supplementary Figure 5: Increased Levels of Saturated VLCFA Species in Third Instar Larvae with Abcd1 LOF**. When lipids were analyzed using LCMSMS, a more pronounced increase in C26:0 levels were observed in the LOF models compared to GCMS (**Fig. 3**). Both RNAi strains demonstrated a 3.3-3.4-fold elevation of C26:0 compared to *yw* and a 3.4-3.6-fold increase compared to da>Luc. The *Abcd1* deletion model (Abcd1^Δ4/Δ4^) exhibited a 3.2-fold rise compared to *yw* and 2.4-fold rise compared to heterozygotes for the deletion (Abcd1^Δ4/+^), the latter accumulated C26:0-canitine more than *yw* by only 1.2. Similarly, elevations were observed for C22:0-carnitine together with C24:0-carnitine and C26:0-carnitine (as a single measurement) in these genotypes compared to control (1.7-3.5-fold for RNAi, 3.6-fold for Abcd1^Δ4/Δ4^). Abcd1^Δ4/Δ4^ showed a 2-fold increase in these three acylcarnitines compared to Abcd1^Δ4/+^. For lysophosphatidylcholine derivatives (LPC, bottom two panels), no significant elevations were detected between LOF and control. The acaylcarnitines were analyzed by LCMSMS (refer to materials and methods for details on multiple t-test comparisons).

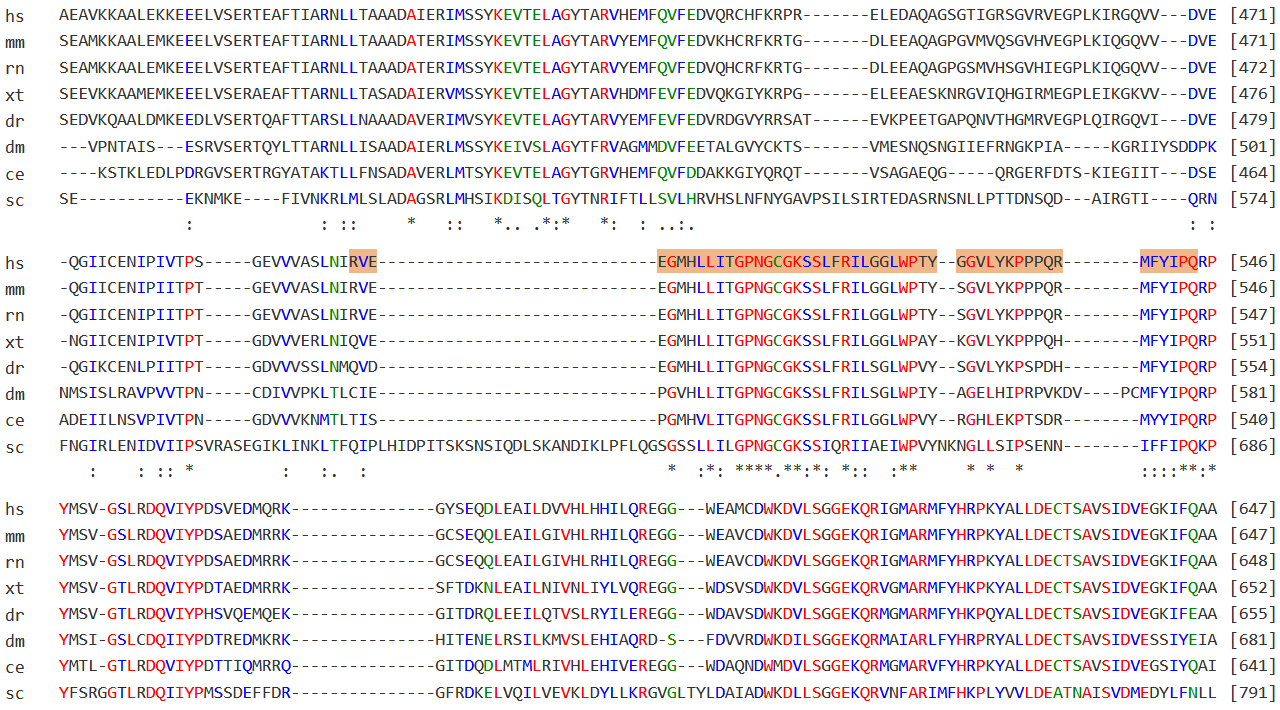

**Supplementary Figure 6: Alignment of ABCD1 Sequences Across Species**. This figure shows an alignment of the amino acid sequence of human ABCD1 (P33897) with various species, covering from exon 3 to exon 9. The segment corresponding to human exon 5 is highlighted in orange, demonstrating near-perfect homology with the species compared here (similarly for the adjacent exons). Red shading highlights the location of the recurrent human variant R518Q, situated well within the homologous region, and perfectly conserved across all the species presented. Abbreviations for species are as follows: hs = Homo sapiens; mm = Mus musculus; rn = Rattus norvegicus; xt = Xenopus tropicalis; dr = Danio rerio; dm = Drosophila melanogaster; ce = Caenorhabditis elegans; sc = Saccharomyces cerevisiae. The image was created using the MARRVEL aggregator (MARRVEL.org).^7^

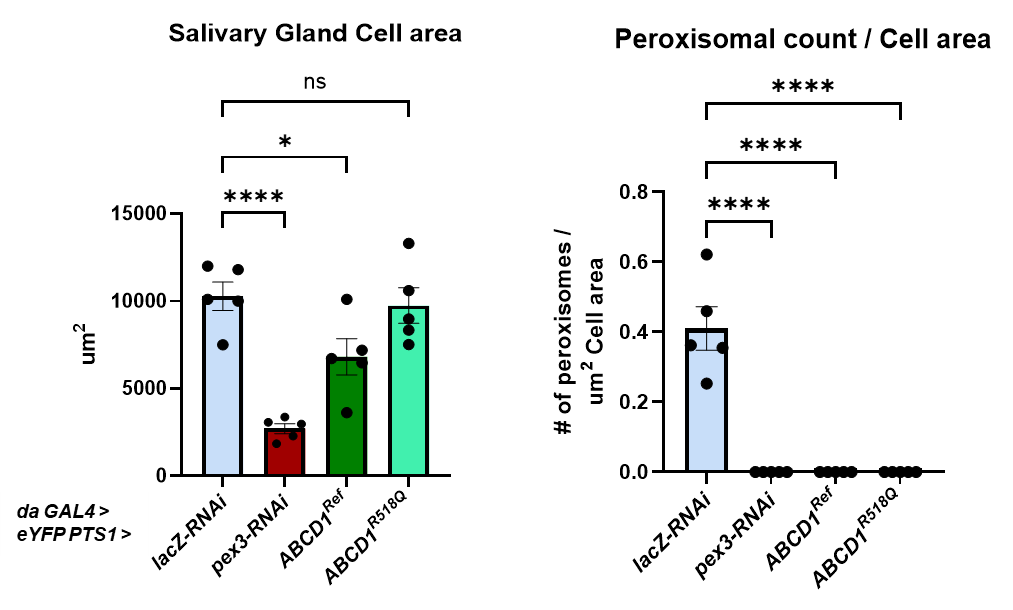

**Supplementary Figure 7: Salivary gland cell area in um^2^.** Human *ABCD1^Reference^* when overexpressed ubiquitously at 29^0^C affects salivary gland cell area as compared to control *lacZ-RNAi* but the variant *ABCD1^R518Q^* fails to do so.

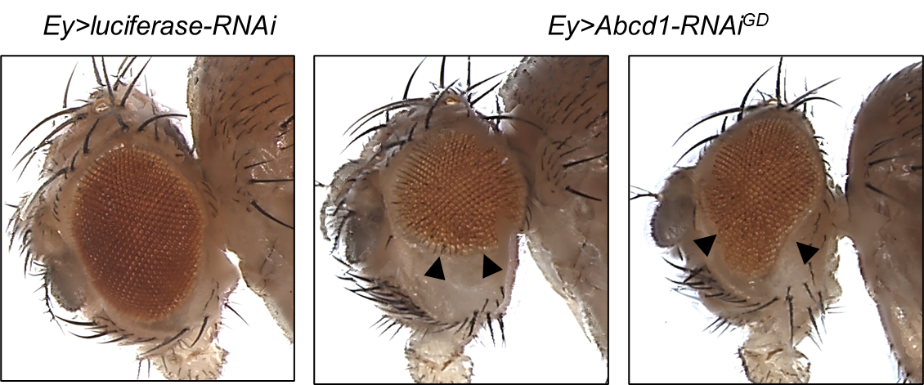

**Supplementary Figure 8**: ***Abcd1*-knockdown with Eyeless-GAL4**. *Ey>Abcd1-RNAi^GD^* flies show abnormal eye development, either small rounded eye or a drop-like shaped eye, with normal pigmentation, in comparison to control *Ey>Luciferase-RNAi.* The penetration of the phenotype is less than 30%.

**SUPPLEMENTARY FILE REFERENCES**

1.	Sivachenko A, Gordon HB, Kimball SS, Gavin EJ, Bonkowsky JL, Letsou A. Neurodegeneration in a Drosophila model of adrenoleukodystrophy: the roles of the Bubblegum and Double bubble acyl-CoA synthetases. Disease models & mechanisms. 2016;9(4):377-87. Epub 2016/02/20. doi: 10.1242/dmm.022244. PubMed PMID: 26893370; PMCID: Pmc4852500.

2.	Gordon HB, Valdez L, Letsou A. Etiology and treatment of adrenoleukodystrophy: new insights from Drosophila. Disease models & mechanisms. 2018;11(6). Epub 2018/05/10. doi: 10.1242/dmm.031286. PubMed PMID: 29739804; PMCID: Pmc6031365.

3.	Pujol A, Hindelang C, Callizot N, Bartsch U, Schachner M, Mandel JL. Late onset neurological phenotype of the X-ALD gene inactivation in mice: a mouse model for adrenomyeloneuropathy. Human molecular genetics. 2002;11(5):499-505. doi: 10.1093/hmg/11.5.499. PubMed PMID: 11875044.

4.	Strachan LR, Stevenson TJ, Freshner B, Keefe MD, Miranda Bowles D, Bonkowsky JL. A zebrafish model of X-linked adrenoleukodystrophy recapitulates key disease features and demonstrates a developmental requirement for abcd1 in oligodendrocyte patterning and myelination. Human molecular genetics. 2017;26(18):3600-14. doi: 10.1093/hmg/ddx249. PubMed PMID: 28911205; PMCID: PMC5886093.

5.	Sayers EW, Bolton EE, Brister JR, Canese K, Chan J, Comeau DC, Connor R, Funk K, Kelly C, Kim S, Madej T, Marchler-Bauer A, Lanczycki C, Lathrop S, Lu Z, Thibaud-Nissen F, Murphy T, Phan L, Skripchenko Y, Tse T, Wang J, Williams R, Trawick BW, Pruitt KD, Sherry ST. Database resources of the national center for biotechnology information. Nucleic acids research. 2022;50(D1):D20-d6. doi: 10.1093/nar/gkab1112. PubMed PMID: 34850941; PMCID: PMC8728269.

6.	Fourcade S, López-Erauskin J, Galino J, Duval C, Naudi A, Jove M, Kemp S, Villarroya F, Ferrer I, Pamplona R, Portero-Otin M, Pujol A. Early oxidative damage underlying neurodegeneration in X-adrenoleukodystrophy. Human molecular genetics. 2008;17(12):1762-73. Epub 2008/03/18. doi: 10.1093/hmg/ddn085. PubMed PMID: 18344354.

7.	Wang J, Al-Ouran R, Hu Y, Kim SY, Wan YW, Wangler MF, Yamamoto S, Chao HT, Comjean A, Mohr SE, Perrimon N, Liu Z, Bellen HJ. MARRVEL: Integration of Human and Model Organism Genetic Resources to Facilitate Functional Annotation of the Human Genome. American journal of human genetics. 2017;100(6):843-53. Epub 20170511. doi: 10.1016/j.ajhg.2017.04.010. PubMed PMID: 28502612; PMCID: PMC5670038.
